# Species-specific differences in the susceptibility of fungi towards the antifungal protein AFP depend on C3 saturation of glycosylceramides

**DOI:** 10.1101/698456

**Authors:** Norman Paege, Dirk Warnecke, Simone Zäuner, Silke Hagen, Ana Rodrigues, Birgit Baumann, Melanie Thiess, Sascha Jung, Vera Meyer

## Abstract

AFP is an antimicrobial peptide (AMP) produced by the filamentous fungus *Aspergillus giganteus* and a very potent inhibitor of fungal growth without affecting the viability of bacteria, plant or mammalian cells. It targets chitin synthesis and causes plasma membrane permeabilization in many human and plant pathogenic fungi, but its exact mode of action is not known. We have recently proposed adoption of the “damage-response framework of microbial pathogenesis” put forward by Pirofksi and Casadevall in 1999 regarding the analysis of interactions between AMPs and microorganisms, thus, predicting that the cytotoxic capacity of a given AMP is relative and depends not only on the presence/absence of its target(s) in the host and the AMP concentration applied but also on other variables, such as microbial survival strategies. We show here using the examples of three filamentous fungi (*Aspergillus niger, Aspergillus fumigatus, Fusarium graminearum*) and two yeasts (*Saccharomyces cerevisiae, Pichia pastoris*) that the important parameters defining the AFP susceptibilities of these fungi are (i) the presence/absence of glycosylceramides, (ii) the presence/absence of Δ3(*E*)-desaturation of the fatty acid chain therein, and (iii) the (dis)ability of these fungi to respond to AFP inhibitory effects with the fortification of their cell walls via increased chitin and β-(1,3)-glucan synthesis. These observations support the adoption of the damage-response framework to holistically understand the outcome of AFP inhibitory effects.

**Importance:** Our data suggest a fundamental role of glycosylceramides in the susceptibility of fungi towards AFP. We discovered that only a minor structural difference in these molecules – the saturation level of their fatty acid chain, controlled by a 2-hydroxy fatty N-acyl-Δ3(*E*)-desaturase – is a key to understanding the inhibitory activity of AFP. As glycosylceramides are important components of fungal plasma membranes, we propose a model which links AFP-mediated inhibition of chitin synthesis in fungi with its potential to disturb plasma membrane integrity.

## Introduction

The continuous rise of plant, animal and human pathogenic fungi, being virtually resistant against the antifungal drugs used currently, calls urgently for novel antifungal substances and new antifungal strategies (1). In addition, most of the antifungal drugs in use exert cytotoxic effects also on humans because the target molecules are not fungal-specific but are also present in higher eukaryotes (2). Novel antifungals with specific targets only present in fungi is, thus, one important prerequisite for the development of new fungal-specific drugs. An interesting lead compound for a new generation of fungicides is the antifungal protein AFP from *Aspergillus giganteus*. This is not only because of its fungal-specific mode of action in the micromolar range, but also because of its high stability against temperature (80 °C), pH (2–10) and proteases (3). It has, furthermore, been demonstrated that treatment of plants used agriculturally with AFP or a transgenic expression of the *afp* gene in these plants can protect them against fungal infections caused by *Blumeria graminis*, *Fusarium oxysporum* and *Magnaporthe grisea* (*Pyricularia grisea*), to name but a few (3–5).

AFP is a small, 51-amino acid, cationic and amphipathic peptide, whose three-dimensional structure has been resolved by NMR (4). It is the founder molecule of the AFP family, which consists of more than 50 members present in about 30 different *Ascomycota* (6). The structural characteristics of this family include a γ-core motif, six conserved cysteine residues and a highly stable beta-barrel folding. Notably, the γ-core is a common feature in all cysteine-stabilized antimicrobial peptides (AMPs) from bacteria, fungi, plants and (in)vertebrates (7). It has recently been demonstrated by *in silico* molecular dynamics simulations that AFP interacts strongly with a fungal membrane model without penetrating it, whereby its γ-core motif was actively involved in the formation of the membrane-AFP binding interface (4). This agrees with electron microscopic data reporting that AFP binds heavily to the cell wall and plasma membrane of sensitive fungi (e.g. *A. niger*) but not to the cell surface of AFP-resistant fungi (e.g. *Penicillium chrysogenum*). In fact, AFP binds to chitin *in vitro* and has been shown to inhibit chitin biosynthesis *in vivo* (8). Chitin is directly adjacent to the plasma membrane and an important structural component of the fungal cell wall together with β-(1,3)-glucan, β-(1,6)-glucan, α-(1,3)-glucan, (galacto)mannans and glycoproteins (9). In addition to its ability to disturb chitin biosynthesis in susceptible fungi, AFP has also been shown to stretch and permeabilize their plasma membranes within minutes after application (8, 10, 11).

Fungal chitin biosynthesis is far from being understood. It is assumed that chitin is synthesized at the plasma membrane by transmembrane-localized chitin synthases (CHSs) that are transported to the plasma membrane in an inactive form within chitosomes (a specific population of secretory vesicles) and become activated after plasma membrane insertion (9). Yeast and filamentous fungal genomes contain several CHS encoding genes (up to 12 per genome) thought to fulfill different functions during growth and developmental processes (12). The three-dimensional structure has been resolved for none of the CHS so far, hence, it is currently impossible to predict or deduce any direct interaction of AFP with a CHS based on structural information. A number of possible scenarios have, thus, been proposed which could explain the inhibitory effect of AFP being localized at the outside of the fungal plasma membrane and exerting effects on both chitin biosynthesis and plasma membrane integrity (3): (i) AFP might prevent the fusion of chitosomes with the plasma membrane; (ii) AFP might disturb the proper embedding of CHSs in the plasma membrane, for example, by interacting with adjacent plasma membrane components; and/or (iii) AFP might interfere with the enzymatic activity of CHSs, for example, by binding to newly synthesized chitin, thus, preventing polymerization of the nascent chain. All three events are conceivable and not mutually exclusive. They would eventually cause membrane stretching due to malformation of the cell wall which can no longer withstand the internal turgor pressure. This, in turn, would lead to cell lysis, predominantly at the hyphal tips where cell wall biosynthesis mainly occurs. In agreement, tip-localized bursting is frequently observed in the highly AFP-susceptible *A. niger* when treated with AFP (8, 10, 11).

Fungi use different survival strategies to fight against any lethal effects of AFP. The analysis of different wild-type and cell wall mutants of *Saccharomyces cerevisiae*, *A. niger* and *Fusarium oxysporum* uncovered the fact that fungi which are less susceptible towards AFP fortify their cell walls with chitin in response to AFP, whereas fungi which are more susceptible towards AFP fail to do so (13). This was mechanistically explained by the observation that less susceptible strains rely on the calcium/calcineurin/Crz1p signaling pathway to reinforce chitin synthesis, whereas susceptible strains deploy the Pkcp/Slt2p/Rlm1p cell wall integrity pathway, whose main output is glucan and not chitin synthesis (13). This observation led us to propose the adoption of the “damage-response framework of microbial pathogenesis” (14–16) to explain the effect of AFP on fungi. When translating the tenets of this conceptual approach to AFP fungal interactions, it can be postulated that the outcome of an AFP attack is dependent on i) the innate susceptibility of the microorganism, ii) the damage potential of AFP defined by its concentration and target-(non)specific molecular interactions, and iii) the survival response, which can be appropriate, too weak or too strong and, thus, detrimental to the host. Consequently, the damage-response framework predicts that the cytotoxic capacity of AFP on fungi is relative (13).

In order to further study the fungal-specific mode of action of AFP, the different susceptibilities of fungi towards AFP and the link between its cell wall and cell membrane effects, we searched the literature for plasma membrane components which are not present in prokaryotes but are in eukaryotes and, most importantly, differ between eukaryotic kingdoms. This is the case for sphingolipids, which are present in eukaryotes but differ in their composition in different kingdoms (17). Very simple forms of sphingolipids are glycosylceramides (GlyCer), which are prevalent in fungi (Fig. 1). They consist of a sphingoid base, a fatty acid and either a glucose (GlcCer) or galactose (GalCer) moiety. Fungal GlyCer can be distinguished from those in plants and mammals by a methyl group branching from C9 of the sphingoid base. In addition, there are variable levels of unsaturation and lengths of the fatty acid chain (Fig. 1, (18)). A further literature survey for data on fungi whose GlyCer composition has been resolved and for which the minimal inhibitory concentration of AFP is known, surprisingly uncovered that the susceptibility of fungi towards AFP seemed to match the presence or absence of an Δ3(*E*)-double bond within the fatty acid (Fig. 1 and Table 1). The highly susceptible *A. niger*, for example, contains unsaturated GlyCer, the less susceptible *A. nidulans* contains saturated GlyCer and the AFP-resistant *S. cerevisiae* does not contain GlyCer at all. Importantly, the enzyme synthesizing the unsaturation at the C3 atom has been studied in *F. graminearum* (2-hydroxy fatty N-acyl-Δ3(*E*)-desaturase) (19). In the present study, we, therefore, experimentally investigated differences in the AFP susceptibility of different fungi (*A. niger, A. fumigatus, F. graminearum, Pichia pastoris* [new name *Komagataella phaffii*]*, S. cerevisiae*) dependent on the presence or absence of GlyCer and their saturation levels. In doing so, we followed knock-out and knock-in approaches of different orthologous genes predicted to encode a Δ3(*E*)-desaturase and additionally performed different chemical assays to inhibit GlyCer biosynthesis in these fungi. The data presented here demonstrate a direct correlation between the un/saturation state of GlyCers and fungal susceptibilities towards AFP.

**Figure 1:**
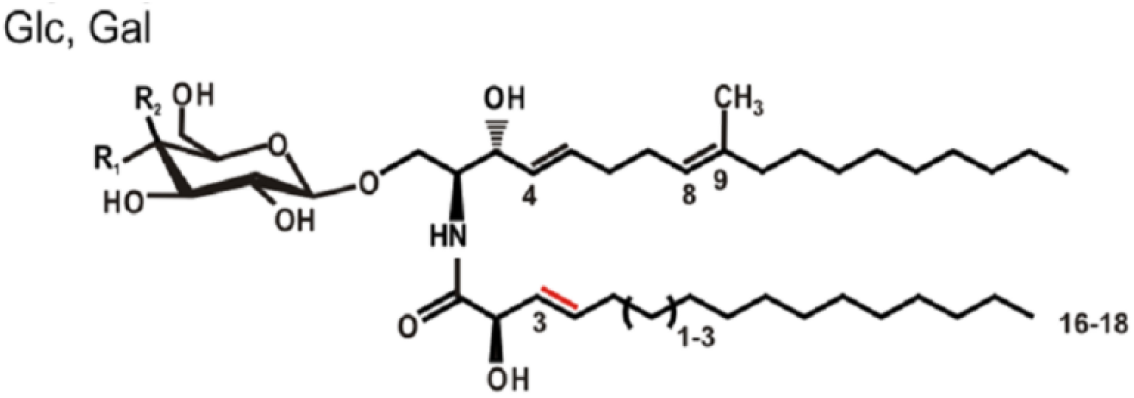
Fungal glycosylceramides consist of either a glucose (Glc) or galactose (Gal) moiety, a sphingoid base and a fatty acid. The double bond generated by the 2-hydroxy fatty N-acyl-Δ3(*E*)-desaturase is marked in red.

**Table 1:**
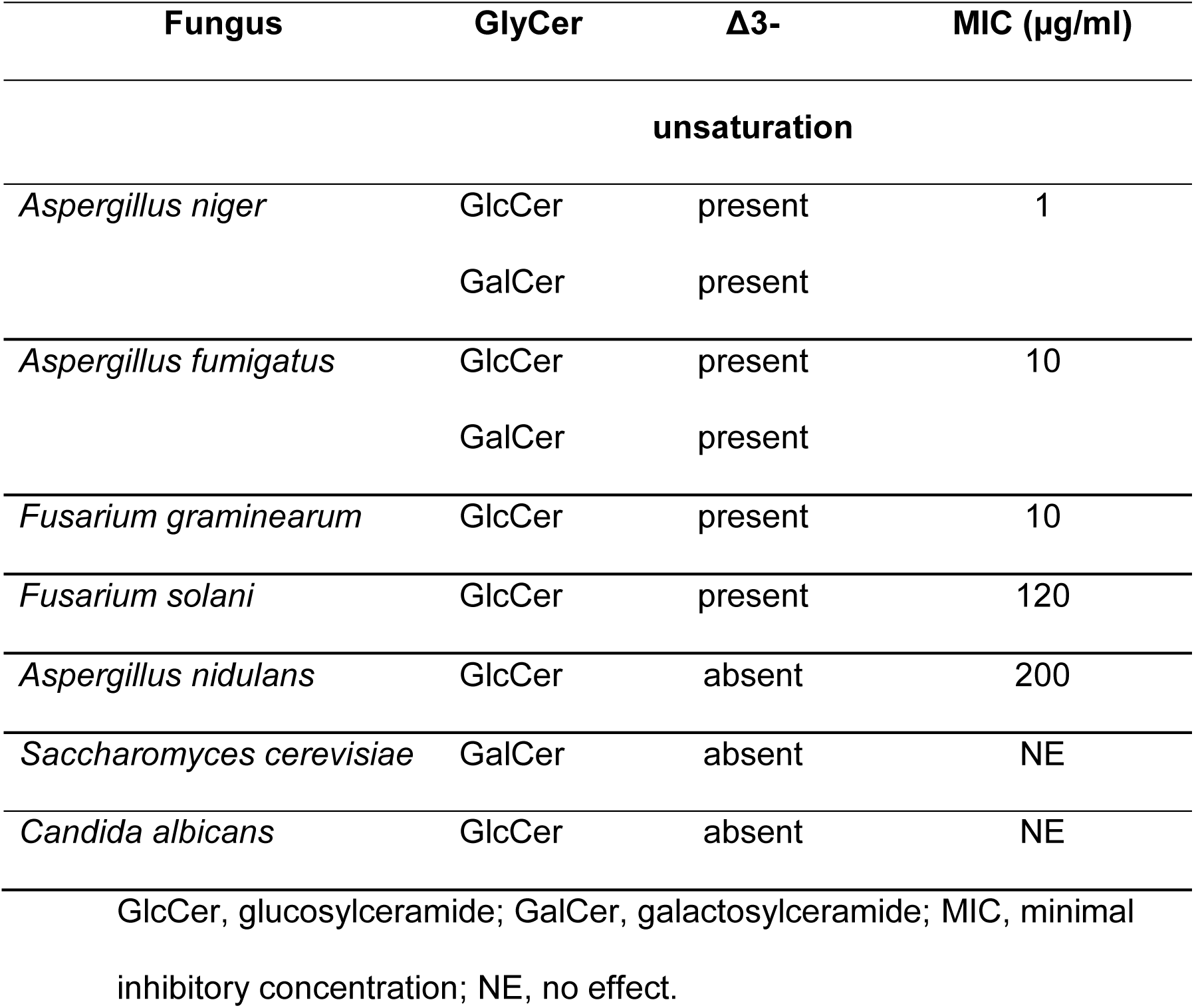
Literature data for fungal glycosylceramides (GlyCer, (18)) and minimal inhibitory concentrations (MIC) of AFP (11).

## Methods

### Strains and cultivation conditions

Fungal strains used in this study are listed in Table 2. *S. cerevisiae* and *P. pastoris* were cultivated in yeast peptone glucose medium (YPG, 0.3% yeast extract, 2% peptone, 1% glucose, 0.5% NaCl), buffered complex glycerol (BMGY) or methanol medium (BMMY) medium (1% yeast extract, 2% peptone, 100 mM potassium phosphate, pH 6.0, 1.34% yeast nitrogen base, 4*10E-5% biotin, 1% glycerol or 0.5% methanol) (20) at 30 °C. *F. graminearum* strains were cultivated in minimal medium (MM) or complete medium (CM, which is MM supplemented with 1% yeast extract and 0.5% casaminoacids) (20) at 28 °C. *A. niger* strains were cultivated at 28 °C, 30 °C or 37 °C in MM or CM (supplemented with 10 mM uridine if required). Transformations of *A. niger*, selection procedures, genomic DNA extractions and diagnostic PCRs were performed using protocols described recently (21). *A. niger* was transformed using the PEG method described in (22). Standard PCRs, general cloning procedures and Southern analyses were carried out according to established protocols (23). The transformation of *F. graminearum* strains was performed according to (19). Yeast strains were transformed via electroporation, as described previously (24, 25). *P. pastoris* protein production was performed in adapted MM (1.34 % YNB, 0.5 % methanol, 4*10E-5 % biotin) at 30 °C. Chemically competent *E. coli* Top10 (Invitrogen™) cells were transformed using an established heat shock procedure (23) with the desired plasmid or ligation mix followed by cultivation at 37 °C on LB medium containing antibiotic.

**Table 2.** Fungal strains used in this study.

| Spezies | Strain<br>name | Description | Reference |
| --- | --- | --- | --- |
| *A. niger* | N402 | *cspA1* | (26) |
| *A. niger* | AB4.1 | *cspA1*, *pyrG*<sup>-</sup> | (26) |
| *A. niger* | NP1.15 | *cspA1*, ΔAn01g09800::*AopyrG*, N402<br>derivative | this work |
| *A. fumigatus* | ATCC<br>9197 | wild-type | (27) |
| *F. graminearum* | 8/1 | wild-type | (28) |
| *F. graminearum* | Δ3FgKO | ΔFJ176923 | (19) |
| *P. pastoris* | GS115 | *his4*<sup>-</sup> | (29) |
| *P. pastoris* | BBA21.3 | GS115 transformed with pNP1.13 | This work |
| *P. pastoris* | NP7.1 | GS115 transformed with pPIC3.5 | This work |
| *P. pastoris* | SZ51 | pGAPZ/C+ FJ176923 | (19) |
| *S. cerevisiae* | BY4741 | MATa *his3*Δ1 *leu2*Δ0 *met15*Δ0 *ura3*Δ0 | (30) |
| *S. cerevisiae* | BBA20.2 | BY4741 transformed with pBBA 5.1 | This work |

### Molecular cloning

In order to clone the cDNA of the predicted Δ3(*E*)-desaturase from *A. niger* (An01g09800), total RNA from a culture of *A. niger* strain N402 was extracted from its biomass following the TRIzol™ method (31) and treated with DNAse using the “Ambion DNA free” Kit (Invitrogen™). cDNA was generated from total RNA using a Thermo Fisher RevertAid H Minus First Strand cDNA Synthesis Kit with an Oligo(dT)_18_ primer, according to the manufacturer’s instructions. An01g09800 cDNA was used as the PCR template for the Q5 proofreading polymerase (NEB). Sequence alignments of the PCR product confirmed the predicted intron-exon boundaries of An01g09800. This PCR product also served as a template for a PCR to amplify an open reading frame fragment for sticky end ligation into plasmid pPIC3.5 (32) restricted with EcoRI and NotI. The resulting plasmid pNP1.13 was linearized with SacI and transformed into the *P. pastoris* strain GS115 (Invitrogen™) via electroporation, giving strain BBA21.3. Total cDNA from the *F. graminearum* strain 8/1 (19) was used as a template for the amplification of the Δ3(*E*)-desaturase gene FJ176923 with Pwo-polymerase (peqlab) and ligated into vector pGEM®-T (Promega). After restriction with EcoRI and NotI, the corresponding fragment containing Δ3(*E*)-desaturase was ligated into pGAPZ/C (Invitrogen™), leading to the plasmid, which, after linearization with BglII, was transformed into the *P. pastoris* strain GS115, resulting in the strain SZ51. The cDNA of the Δ3(*E*)-desaturase gene An01g09800 was further used as a template to amplify a fragment for Gibson assembly with fragments of the *S. cerevisiae* TEF promoter and CYC terminator into the vector pYIP5 (Addgene) restricted with HindIII and SalI. PCR amplification of the cassette via Q5 polymerase for sticky-end cloning into vector pRS306 (Addgene) after restriction with the enzymes XhoI and XbaI led to plasmid pBBA5.1, which was transformed into *S. cerevisiae* strain BY4741 (33) via electroporation. The resulting strain was named BBA20.2. All DNA constructs were verified by sequencing before transformation. qPCR was carried out using “Maxima sybr green/ROX qPCR Master Mix” (Thermo Fisher), following the manufacturer’s instructions. Up to 500 ng cDNA was used as a template. The actin gene served as a control (34).

### Identification and deletion of the *A. niger* Δ3(*E*)-desaturase gene

The Δ3(*E*)-desaturase sequence of *F. graminearum* FJ176922.1 (19) was used as a reference to identify the orthologue in *A. niger* via pBLAST (35). The identified open reading frame An01g09800 displayed a sequence identity of 61 % on the amino acid level to FJ176922.1. We used the split marker approach (36) with about 500 ng gel-purified PCR product each to replace An01g09800 with p*yrG* via a crossing-over event. The forward bipartite was created using primers 677 and 462 (see Suppl. Table S1) with the upstream region of *An01g09800* and *pyrG* as a template. The reverse bipartite was amplified with primers 461 and 680 (Suppl. Table S1) using *pyrG* and the *An01g09800* downstream fragment as a template. After purification on MM twice, genomic DNA of the transformants was tested via PCR using primers 677 and 716 (Suppl. Table S1). Southern blot analysis using SalI for the fragmentation of genomic DNA and a digoxigenin-labelled DNA probe binding fragments containing *pyrG* confirmed deletion of *An01g09800* in two strains which we named NP1.15 and NP1.16, respectively.

### Purification and analysis of GlyCer

The biomass of the *A. niger* and *P. pastoris* strains were frozen in liquid nitrogen, freeze-dried and weighted. An amount of 0.7 g dry biomass was ground and extracted twice with 20 ml chloroform and methanol (2:1 v/v), vortexed with glass beads and gently shaken for 2 h. A quantity of 1 ml demineralized water was added before centrifugation (340 g, 4 min). The supernatant was mixed with 7 ml of 0.75 % NaCl solution in water and centrifuged before the bottom phase was separated, and the organic phase was evaporated at 50 °C covered by argon as an inert gas. The sample was resuspended in 1–3 ml chloroform and methanol (2:1 v/v) and filtered through a cotton-plugged Pasteur pipette. Samples, each 5 µl, were analyzed via thin-layer chromatography (TLC) using silica gel plates and a runtime of 1 h in chloroform and methanol (85:15, v/v). The GlcCer from *P. pastoris* was applied as a standard. Visualization of the samples via 8-anilinonaphthalene-1-sulfonic acid under UV light followed. The final detection was carried out with alpha-naphthol sulfuric acid and heating to 170 °C. The GlyCer-positive samples were used for subsequent preparative TLC to purify the GlyCer from the lipids extracted. Bands containing the GlyCer were scratched from the TLC plates and suspended in 6 ml chloroform and methanol (2:1 v/v), before 1.5 ml of 0.75 % NaCl in water was added. After the centrifugation and separation of the organic phases, the samples were filtered through cotton-plugged Pasteur pipettes until all remaining solids had been removed and the liquid was evaporated at 50 °C using argon gas. Samples were reconstituted in 100 µl chloroform and methanol (2:1 v/v) and stored at −20 °C until they were analyzed.

### Mass spectrometry of GlyCer

An amount of 1 µl of purified GlyCer solution or peptide calibration standard (8206195, Bruker Daltonics) were sandwiched in 0.5 µl α-cyano-II-hydroxycinnamic acid on a MALDI ground steel target (Bruker). The acquisition of MALDI spectra was performed using the ultraflex III^™^ mass spectrometer (Bruker Daltonics) in reflector mode detecting positively charged ions in the range of 400-4000 *m/z*. Each sample was analyzed for the expected signals at 778 m/z and 776 m/z for saturated GlyCer and unsaturated GlyCer, respectively. In addition, the instrument was operated in MS/MS mode for the parental ion masses 777 and 778 as mentioned before. The evaluation of the spectra was carried out using the program “mMass - Open Source Mass Spectrometry Tool” (37).

### Chitin and β-1,3-glucan quantification

*P. pastoris* strains were inoculated from preculture grown in BMMY medium for 24 h and incubated in fresh BMMY medium for 48 h. The methanol was supplemented every 24 h to induce the expression of the Δ3(*E*)-desaturase. An amount of 50 µg/ml AFP were added at the start of the cultivation when needed. 10E6 spores/ml from *A. niger* N402 (wild-type) were cultivated in 100 ml YPG medium at 28 °C for 24 h. An amount of 20 µg/ml AFP was added and the cultivation continued up to 72 h. Biomass was harvested via centrifugation (3090 g, 10min) or by filtration (Sartorius, format 3 hw), freeze-dried and used directly for the analysis of β-1,3-glucan content according to (38), with the exception that an ultrasonic bath was used for 1 h instead of ultrasonic treatment for 30 s. The culture supernatant was discarded. The chitin content was measured according to (39). In brief, 50 mg freeze-dried biomass was used for the cell wall extraction. An amount of 4 mg of freeze-dried cell wall fraction was used for the chitin extraction. Calibration curves were generated with curdlan for β-1,3-glucan and glucosamine for chitin. Purified β-1,3-glucan or chitin was quantified using fluorescence spectroscopy (excitation 392 nm, emission 502 nm) or photometry at 520 nm.

### Susceptibility assays

The AFP susceptibility assay was performed as described in (8) and (13) to determine the minimal inhibitory concentration. Susceptibility towards AFP in the presence or absence of 35 µM D-threo-1-phenyl-2-decanoylamino-3-morpholino-1-propanol (D-PDMP dissolved in Milli-Q-water) was determined in the same way.

## Results

### Fungal glucosylceramides mediate sensitivity to AFP

The glucosylceramide biosynthetic pathway has been shown to be important for spore germination and hyphal growth in *A. nidulans, A. fumigatus* and *F. graminearum* (40–42). A central position in this pathway is adopted by the ceramide synthase GCS, which transfers a glucose moiety from uridine 5-diphosphate (UDP)-glucose onto the C1 hydroxyl group of the ceramide. We chemically inhibited GCS activity in the AFP-susceptible strains *A. niger* and *A. fumigatus* using D-PDMP to verify the role of GlcCers in the susceptibility of filamentous fungi towards AFP. This compound is a synthetic analogue of ceramide which acts as an antimetabolite and inhibits the covalent bonding of ceramides with glucose. This inhibition, in turn, leads to the absence of GlcCer in filamentous fungi, as shown for *A. nidulans* (43). As depicted in Figure 2, the absence of GlcCer considerably decreased the susceptibility of both test strains towards AFP as the survival rates increased substantially (> 3-fold). However, the addition of D-PDMP did not completely rescue the strains and its protective effect was stronger for *A. niger* than *A. fumigatus*, potentially pointing at strain-specific differences in innate susceptibilities and/or survival responses. Note that the protective effect of D-PDMP decreased with the increasing AFP concentration, implying that GlcCers might not be the only targets of AFP.

**Figure 2.**
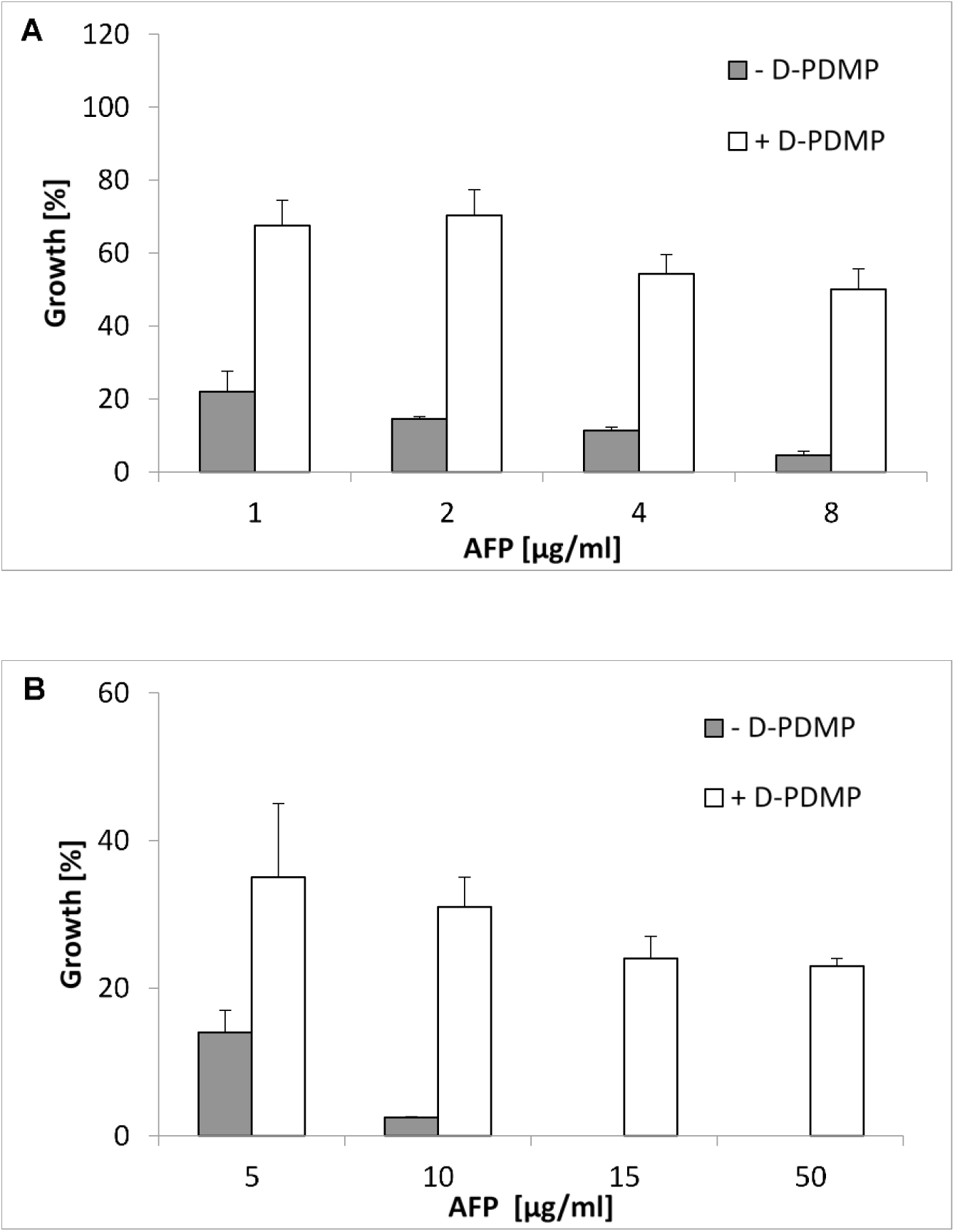
The AFP susceptibility of *A. niger* and *A. fumigatus* is dependent on glucosylceramides. The growth of A) *A. niger* wild-type strain N402 and B) *A. fumigatus* wild-type strain ATCC 9197 at different AFP concentrations in the presence (white bars) or absence (grey bars) of the GlcCer synthesis inhibitor D-PDMP is shown. Data were normalized to control cultures which were not treated with AFP. Data are mean values referring to two biological replicates, each measured as a technical triplicate. Error bars express standard deviations.

### Δ3(*E*)**-**desaturase gene expression mediates sensitivity to AFP

As summarized in Table 1, we hypothesized that a causal relationship between the Δ3(*E*)-desaturation of GlyCer and the susceptibility of fungi towards AFP might exist. In order to study this in detail, we wanted to identify the gene encoding the Δ3(*E*)-desaturase in the AFP-susceptible *A. niger* to generate a knock-out strain for comparative susceptibility assays.

Complementary to this approach, we wanted to establish knock-in strains in otherwise AFP-resilient strains to test whether the introduction of a gene encoding the Δ3(*E*)-desaturase can switch resistance into susceptibility.

BLAST analyses using the known Δ3(*E*)-desaturase gene from *F. graminearum* (ORF code FJ176922.1) identified only one orthologous gene in *A. niger* which showed a sequence identity of 61 % at the amino acid level (ORF code An01g09800). For brevity, we named this gene *dtdA* (<u>d</u>elta three <u>d</u>esaturase). Analysis of our in-house *A. niger* transcriptomic database, containing expression data for ∼ 14,000 *A. niger* genes under 155 different cultivation conditions, found that *dtdA* is expressed under nearly all cultivation conditions, albeit at very low levels similar to regulatory Rho-GTPases ((44), Suppl. Fig. S1.).

We, thus, generated a *dtdA* deletion strain in *A. niger* (strain NP1.15) and compared its susceptibility to its wild-type strain (strain N402). In addition, we performed an analogous comparison with a wild-type strain of *F. graminearum* (strain 8/1) and its Δ3(*E*)-desaturase deletion derivative (strain Δ3FgKO), which had been established previously (19). As depicted in Figure 3, both deletion strains displayed substantially improved growth in the presence of AFP when compared to their parental strains, demonstrating that both strains are less susceptible towards AFP when their Δ3(*E*)-desaturase genes are inactive.

**Figure 3.**
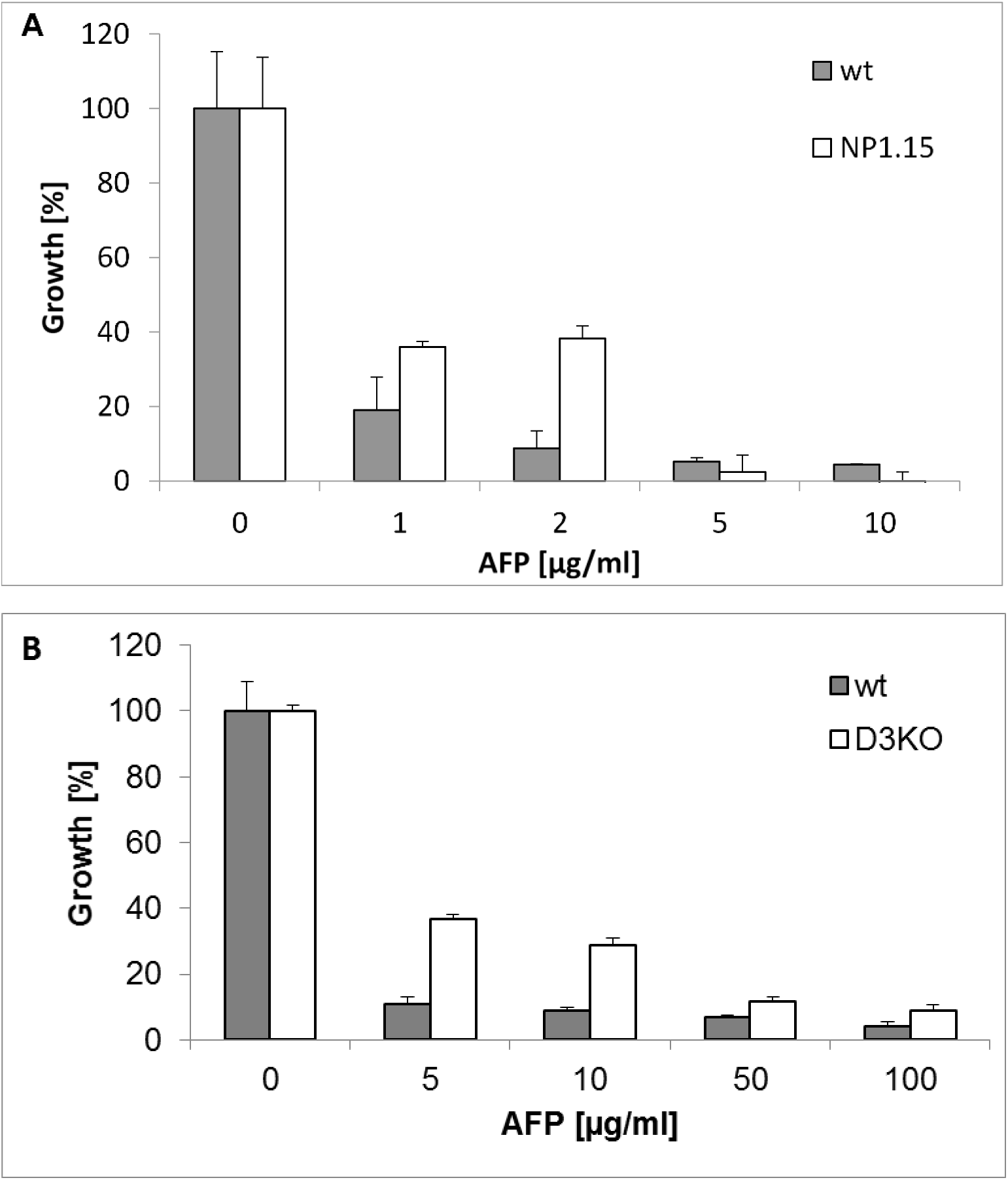
*A. niger* and *F. graminearum* deleted for the Δ3(*E*)-desaturase encoding gene show reduced AFP susceptibilities. (A) Growth of *A. niger* wild-type (wt) strain N402 (grey bars) and Δ*dtdA* strain NP1.15 (white bars) at different AFP concentrations; (B) Growth of *F. graminearum* wild-type (wt) strain (grey bars) and Δ3(*E*)-desaturase knock-out strain (D3KO; white bars). Data were normalized to the growth, i.e. 100 % growth, of control samples not treated with AFP. Data are from two independent experiments, each performed as a technical triplicate. Error bars express standard deviations.

We next put the Δ3(*E*)-desaturase genes from *A. niger* and *F. oxysporum*, respectively, under control of a methanol-inducible promoter and transformed the respective expression constructs into the AFP-resistant yeast *P. pastoris*, giving strain BBA21.3 (expressing the *A. niger* Δ3(*E*)-desaturase gene) and strain SZ51 (expressing the *F. oxysporum* Δ3(*E*)-desaturase gene). Both strains became susceptible to high AFP concentrations when the *dtdA* was expressed, which was not the case for non-inducing cultivation conditions: Strain BBA21.3 (SZ51) displayed 40 % (25%) reduced growth in the presence of 100 µg/ml AFP when forced to express the Δ3(*E*)-desaturase gene. The presence (absence) of Δ3(*E*)-desaturase gene transcripts under inducing (non-inducing) cultivation conditions was proven by qRT-PCR (Suppl. Fig. S2 and data not shown).

A further experiment verified that both the presence of GlyCer and expression of the Δ3(*E*)-desaturase gene are fundamental prerequisites for fungal AFP susceptibility. As *S. cerevisiae* lacks GlyCer (19), one would expect that expression of a Δ3(*E*)-desaturase gene in this AFP-resistant strain would not make it AFP-susceptible, because the substrate for the enzyme is not synthetized by *S. cerevisiae*. Indeed, when the *dtdA* gene of *A. niger* was constitutively expressed in *S. cerevisiae* (BBA20.2), the strain did not show any reduced growth in the presence of 100 µg/ml AFP, despite high mRNA transcript levels of the *dtdA* gene (Suppl. Fig. S3 and data not shown).

### DtdA mediates desaturation of GlyCer fatty acid chains

Mass spectrometry (MS) analyses proved that the *dtdA* gene of *A. niger* encodes a protein exerting an Δ3(*E*)-desaturase enzymatic activity. The GlyCers were isolated and purified by TLC from the *A. niger* wild-type and its *dtdA* deletion strain and from the *P. pastoris* wild-type and its Δ3(*E*)-desaturase knock-in derivatives, the latter cultivated under inducing and non-inducing conditions (Fig. 4 and Table 3). The unsaturation of the fatty acyl moiety of GlyCer was detected indirectly by measurement of the total mass of the purified GlyCers. The mass of the molecular ion [M + Na^+^] 776 m/z corresponds to Δ3(*E*)-unsaturated GlyCer, *N*-(*R*)-2’-hydroxy-(3*E*)-octadec-3-enoyl-1-*O*-β-D-hexosyl-(4*E*,8*E*)-9-methyl-sphinga-4,8-dienine (C43H79O9N), whereas the [M + Na^+^] of the Δ3-saturated counterpart is 778 m/z. Subsequent MSMS analysis led to the fragmentation of the parental ions into the hexosyl-sphingobase fragment [M + Na^+^] 496 m/z and a dimer of the sphingobase fragment [M-2H_2_O + H^+^] 587 m/z. Neither fragment was affected by Δ3(*E*)-desaturation and, therefore, showed a constant molecular mass, regardless of their parental ions of [M + Na^+^] 776 m/z and 778 m/z, respectively. Thus, the mass difference of the parental ions can only be explained by mass differences of the fatty acid moieties due to their unsaturation state. As summarized in Table 3 and Suppl. Fig S4, all fungal strains lacking or not expressing a Δ3(*E*)-desaturase gene showed the [M + Na^+^] 778 m/z signal, which reflects the presence of the double bond in the fatty acid moiety in GlyCer. By contrast, strains expressing a Δ3(*E*)-desaturase gene displayed the [M + Na^+^] 776 m/z signal. Notably, expression of the Δ3(*E*)-desaturase genes into *P. pastoris* did not reach 100 % unsaturation levels. The desaturation efficiency of the *A. niger dtdA* gene reached about 20 % unsaturation level, whereas the desaturation efficiency was 50–100 % for its ortholog from *F. graminearum* (19).

**Figure 4.**
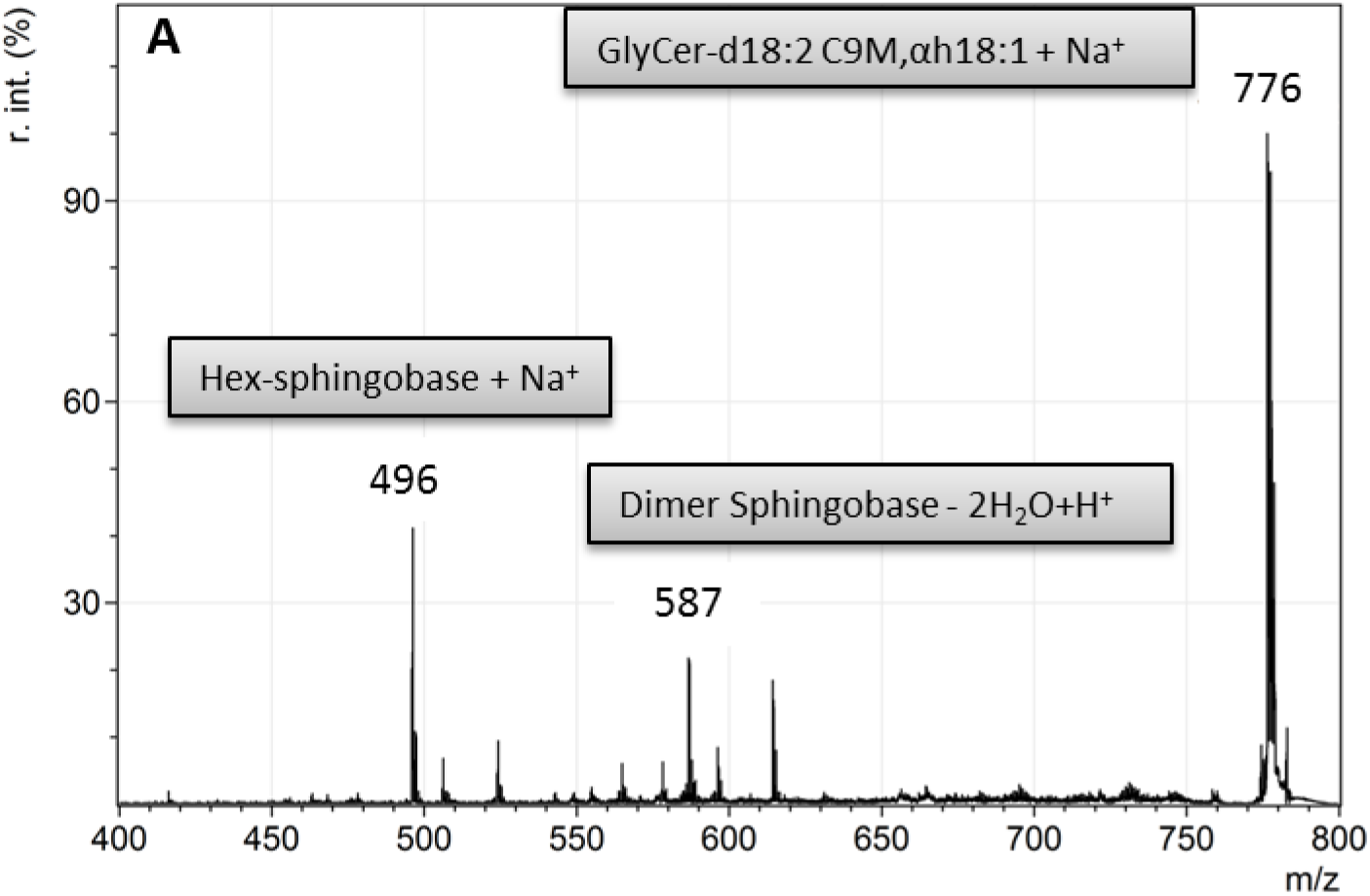
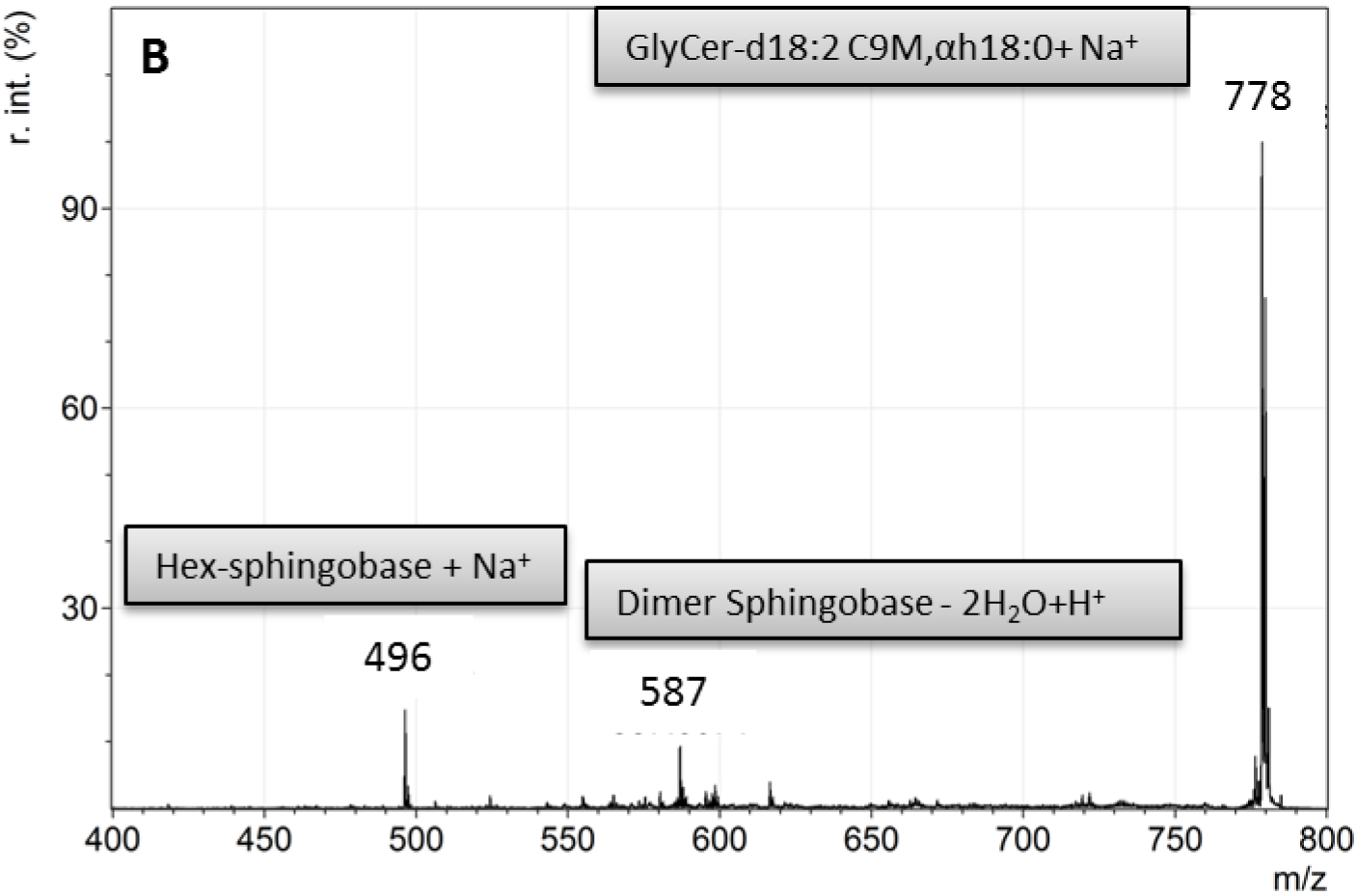
MSMS analysis of *A. niger* wild-type and its *dtdA* deletion strain. Both panels show an overlay of MS spectra for detection of the parental GlyCer ions and the fragmented parental ion masses. (A) Wild-type strain N402. R. int. (%) = relative Intensity in %; (B) *dtdA* deletion strain NP1.15. Hex = Hexose

**Table 3.** MSMS analysis of fungal strains expressing or lacking a Δ3(*E*)-desaturase gene.

| Strain | $\Delta 3(E)$-<br>desaturase | m/z GlyCer<br>[M + Na<sup>+</sup>] | m/z Hexose-<br>sphingobase<br>fragment<br>[M + Na<sup>+</sup>] |
| --- | --- | --- | --- |
| *P. pastoris* BBA21.3 (induced) | Present | 778 (20 %) | 496 |
|  |  | 776 (80 %) | 496 |
| *P. pastoris* BBA21.3 (not induced) | Absent | 778 | 496 |
| *P. pastoris* NP7.1 (not induced) | Absent | 778 | 496 |
| *P. pastoris* NP7.1 (induced) | Absent | 778 | 496 |
| *A. niger* NP1.15 ($\Delta dtdA$) | Absent | 778 | 496 |
| *A. niger* N402 (18 °C) | Present | 776 | 496 |
| *A. niger* N402 (30 °C) | Present | 776 | 496 |
| *A. niger* N402 (37 °C) | Present | 776 (80 %) | 496 |
|  |  | 778 (20 %) |  |

m/z, mass to charge ratio

Because the enzyme activity of the *F. graminearum* Δ3(*E*)-desaturase has been described as temperature-dependent with only 20 % enzyme activity at 30 °C but 100 % at 37 °C (19), we tested whether this is also the case for DtdA. *A. niger* wild-type strain N402 was, thus, cultivated at 18, 30 and 37 °C and *dtdA* transcript levels analyzed by qRT-PCR (Suppl. Fig. S2) and MSMS data recorded for GlyCer populations purified from these biomass samples. As summarized in Table 3, DtdA displays 100 % activity at 30 °C and about 80 % at 37 °C, demonstrating that its temperature-dependent activity is less pronounced and even reciprocal compared to its *F. graminearum* ortholog.

### DtdA prevents strong cell wall reinforcement in *A. niger* and *P. pastoris* in response to AFP

Deletion of *dtdA* did not have any obvious consequences for hyphal growth, biomass accumulation or sporulation of *A. niger* when cultivated in liquid or on solid medium (Fig. 5 and data not shown). Similarly, the presence or absence of the *dtdA* gene did not affect the levels of chitin and β-1,3-glucan in the cell wall of *A. niger* (Suppl. Fig. S5), suggesting that the cellular role of DtdA for growth is rather negligible.

**Figure 5.**
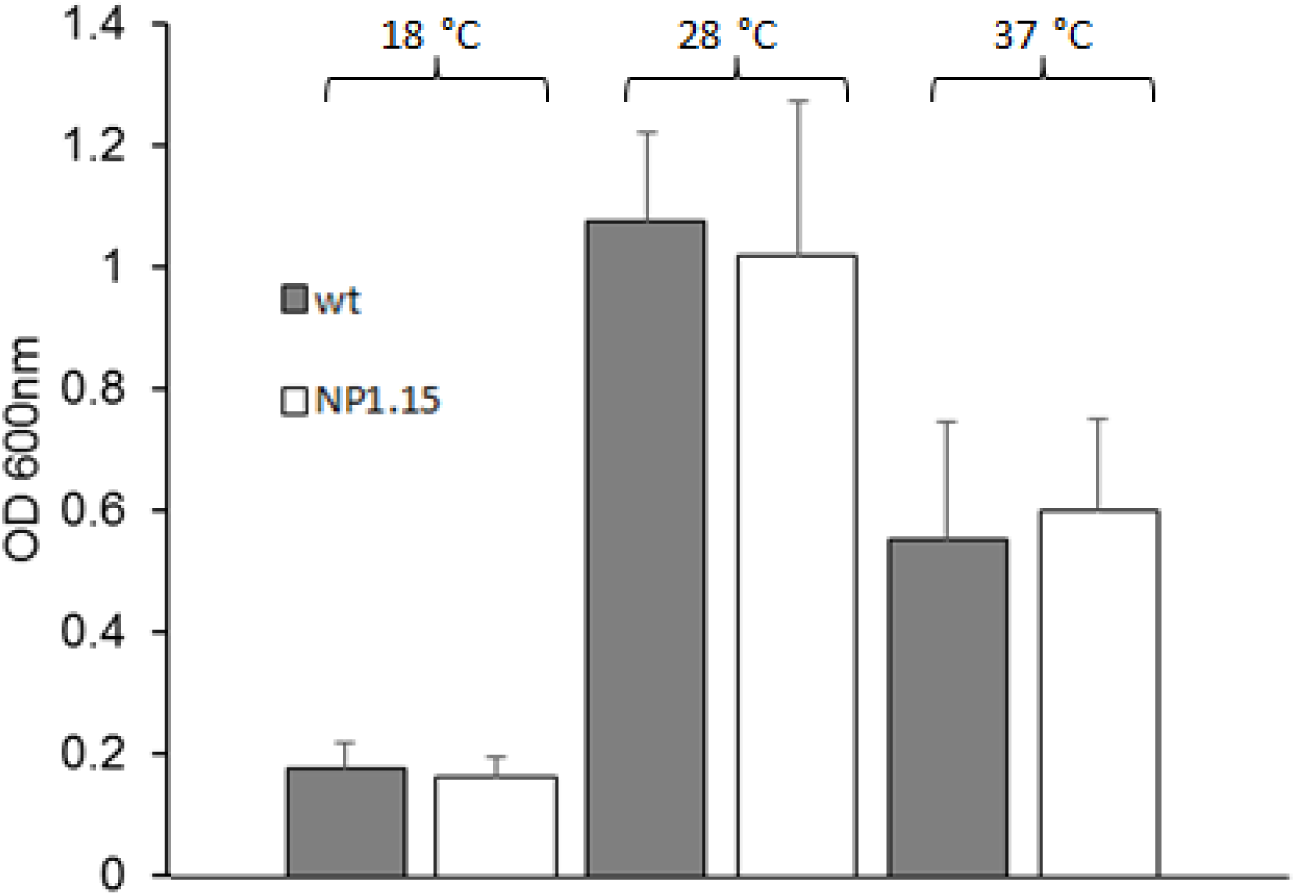
Biomass accumulation of *A. niger* wild-type N402 and its *dtdA* deletion strain. Cultures of *A. niger* wild-type (wt) strain N402 and Δ*dtdA* strain NP1.15 (white bars) were incubated, according to (8), in microtiter plates at three different temperatures for 48 h. Growth was evaluated by determining the optical density (OD) at 600 nm. Data were derived from three independent experiments each performed as a technical triplicate. Error bars express standard deviations.

However, the cell wall stress response of *A. niger* to the presence of AFP differed significantly between the wild-type strain N402 and its *dtdA* deletion derivative, strain NP1.15. Whereas the AFP-susceptible N402 responded to AFP-provoked cell wall stress with a decrease in chitin (−13 %) and an increase in β-1,3-glucan (+26 %) levels, the less AFP-susceptible *ΔdtdA* strain was able to reinforce its cell wall with both chitin (+16 %) and β-1,3-glucan (+78 %) upon AFP treatment (Fig. 6A). Hence, there is a positive correlation between the reduced AFP susceptibility and increased synthesis of cell wall chitin and β-1,3-glucan in *A. niger* upon AFP treatment. A slightly different response was observed for *P. pastoris*. The AFP-resistant wild-type strain responded to the presence of AFP with reduced chitin (−10 %) but increased β-1,3-glucan synthesis considerably (+46 %), whereas the *P. pastoris* strain BBA21.3 expressing the *dtdA* gene and being, thus, AFP-susceptible responded with increased chitin (+6 %) and somehow only a limited increase in β-1,3-glucan synthesis (+20 %) to AFP (Fig. 6C). These observations suggest that (i) both *A. niger* and *P. pastoris* wild types differ in the strength of their AFP survival responses, (ii) the defense mechanism against AFP is strong cell wall fortification with both chitin and β-1,3-glucans, and (iii) that this response is somehow hindered by DtdA and/or the unsaturation level of GlyCers.

**Figure 6.**
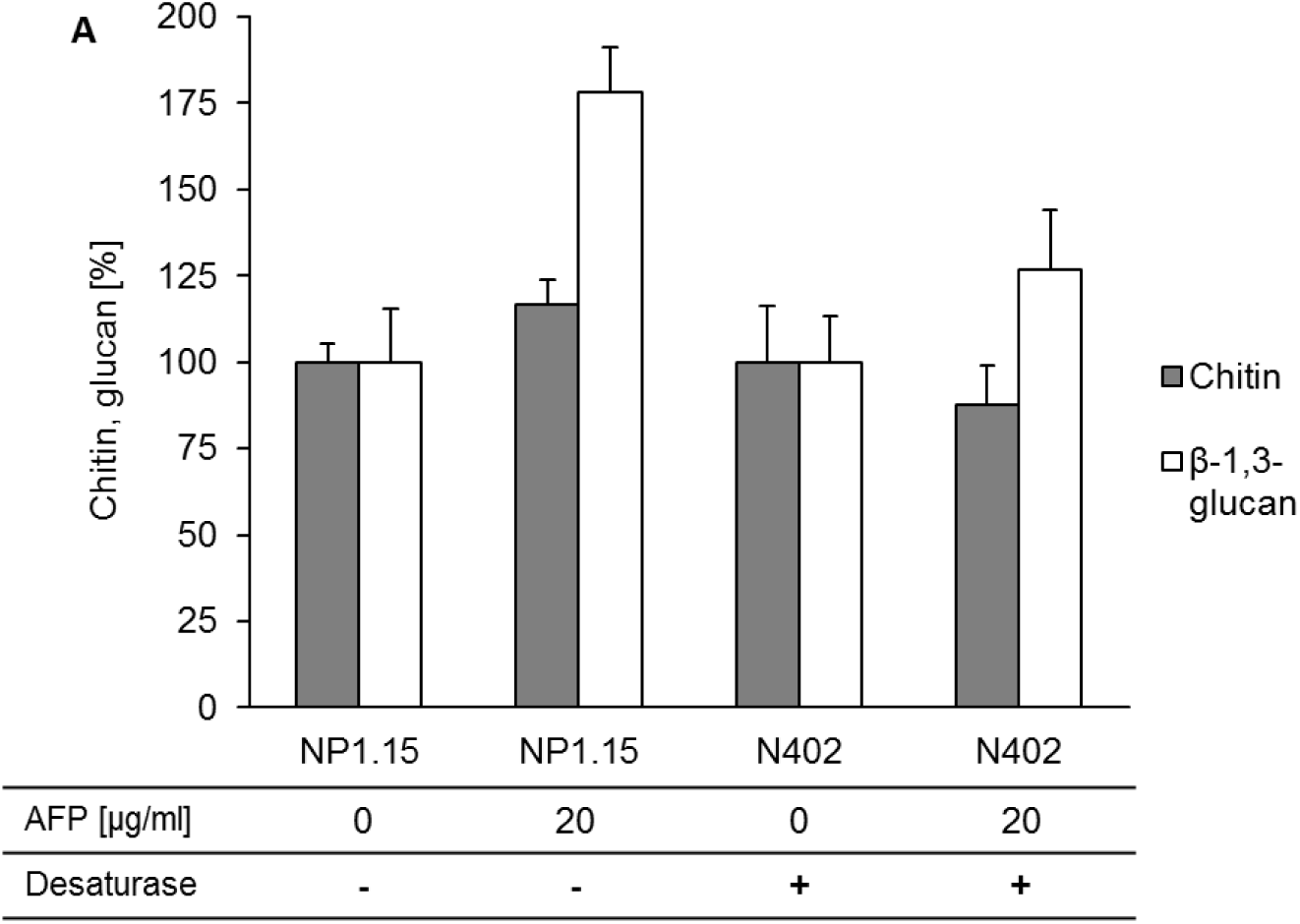
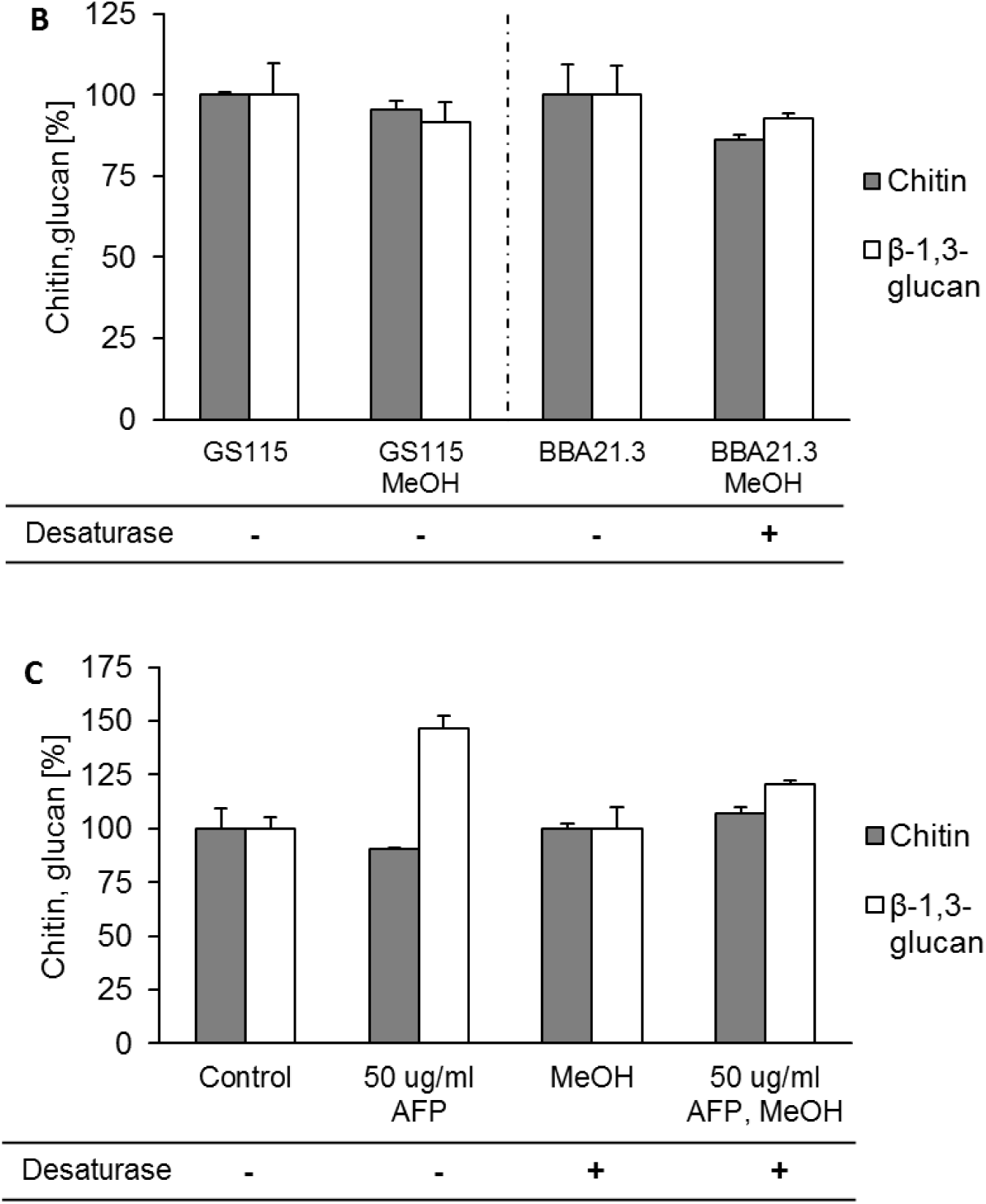
Chitin and β-1,3-glucan responses of *A. niger* and *P. pastoris* when stressed with AFP. The relative amounts of chitin and β-1,3-glucan in % related to control strains, which were set to 100 %, are shown. (A) *A. niger* wild-type N402 and *dtdA* deletion strain NP1.15 in the presence or absence of 20 µg/ml AFP. (B) *P. pastoris* wild-type GS115 and *dtdA* knock-in strain BBA21.3 in the presence or absence of methanol as an inductor of *dtdA* expression. This experiment served as a control to prove that both *Pichia* strains show comparable cell wall compound synthesis in the presence of methanol. (C) *P. pastoris dtdA* knock-in strain BBA21.3 in the presence and absence of AFP and methanol as an inductor of *dtdA* expression. Data are expressed as mean from two independent biological replicates each performed as a technical triplicate. Error bars express standard deviations. Desaturase (+) = *dtdA* expressed; Desaturase (-) = *dtdA* not expressed.

### Both absence of GlyCer and *dtdA* make *A. niger* considerably less vulnerable to AFP

As has been mentioned above, the ceramide synthase inhibitor D-PDMP prevents synthesis of GlcCer, whereas both GlcCer and GalCer are potential targets of DtdA. We, thus, analyzed the consequences of depletion of GlcCer by D-PDMP in the *dtdA* deletion strain devoid of unsaturated GlyCer for the ability of *A. niger* to survive AFP. The *dtdA* deletion strain NP1.15, therefore, was treated with D-PDMP, as described above, and the growth was assessed in the presence of rising AFP concentrations. As depicted in Figure 7, the survival of NP1.15 improved considerably by 15 % (4 µg/ml AFP) up to 36 % (8 µg/ml AFP) when compared to the wild-type strain N402 (Fig. 2A), suggesting that the unsaturation level of both glycosylceramides, GlcCer and GalCer, define the susceptibility of *A. niger* to AFP to a large extent.

**Figure 7.**
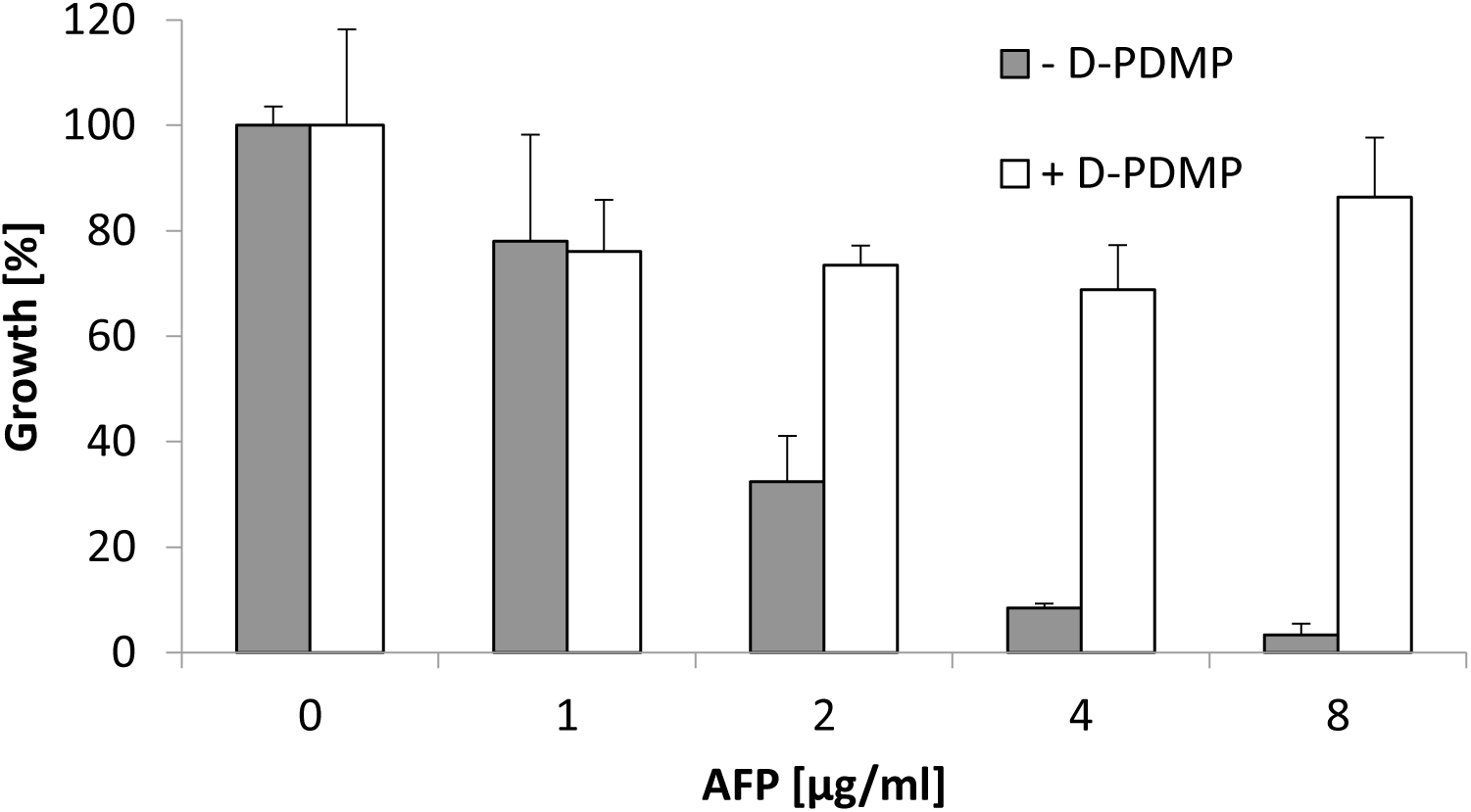
GlyCer-dependent susceptibility of *A. niger dtdA* deletion strain towards AFP. The growth of *A. niger ΔdtdA* strain NP1.15 at different AFP concentrations and in the presence (white bars) or absence (grey bars) of the D-PDMP. Data were normalized to the growth, i.e. 100 % growth, of control samples which were not treated with AFP. NP1.15 spores were treated with the D-PDMP solvent (water) or D-PDMP. Data are expressed as mean from two independent biological experiments performed in a technical triplicate. Error bars express standard deviations.

## Discussion

This is the first study reporting that fungal GlyCers are key molecules defining the susceptibility of fungi towards AFP and that the (un)saturation level of their fatty acid moiety plays a vital role in this relationship. We have interrogated this phenomenon in five different uni- and multicellular fungi and identified a 2-hydroxy fatty N-acyl-Δ3(*E*)-desaturase as the enzyme responsible for the GlyCer desaturation. We named this enzyme DtdA in *A. niger*. Our genetic, chemical and bioanalytical data show conclusively that the presence (or absence) of GlcCer makes a fungus AFP-sensitive (or AFP-insensitive) and that the presence (or absence) of DtdA makes a fungus highly (or lowly) susceptible towards AFP. We, furthermore, provide data which suggest two mechanistic effects to interpret this phenomenon, both of which are not mutually exclusive. Instead, both are in support of a damage-response framework which we proposed earlier (13) and which is neither AFP- nor host-centered: (i) unsaturated GlyCers might be (direct or indirect) targets of AFP (see Figs. 2 and 7) and (ii) saturated GlyCers are a necessary precondition for the proper fortification of fungal cell walls with both chitin and glucans in response to AFP (see Fig. 6). We have shown, using the examples of *S. cerevisiae*, *A. niger* and *P. pastoris*, that they differ in the availability of AFP targets (see Tables 1 and 3) and that *A. niger* and *P. pastoris* also differ in the strength of their cell wall responses to AFP (see Fig. 6). As nothing is known so far about the overall interplay of (un)saturated GlyCers with AMPs and cell wall synthesizing enzymes, the molecular events have remained elusive and need to be scrutinized in the future.

GlyCers are synthesized in the endoplasmic reticulum (ER) and Golgi (46, 47) by ER- or Golgi-resident biosynthetic enzymes and become transported in secretory vesicles to the outer leaflet of plasma membranes (46). In mammals, they are known to modulate cell signaling, including calcium signaling and endocytic trafficking (48, 49). In fungi, it has been shown that they differ between various species (50) and that they are important for polarized growth (51), conidiation (41) and stress resistance (52) (for reviews see <u>53-55</u>). Notably, GlyCer levels are quite low compared to other membrane lipids and reach only about 0.007 % of total lipids in mammalian spleen tissues (56).

Although controversially discussed, GlyCers are thought to form, together with other proteins, special membrane domains called lipid rafts or ceramide platforms because of their different biophysical properties (57, 58). The co-localization of fungal chitin and glucan synthases within lipid rafts has also been documented for the oomycete *Saprolegnia monoica* (59). In view of these observations and the knowledge accumulated for AFP so far (see the Introduction and this study), we propose the following working model (Fig. 8), which will be studied further in future experiments. We hypothesize that AFP accumulates on the surface of the outer leaflet of fungal plasma membranes, as predicted by recent molecular dynamics modeling approaches (4), which confirmed electron and light microscopic data published earlier (8, 10, 11). AFP interacts via its γ-core motif either directly or indirectly with (un)saturated GlyCers embedded in the outer leaflet. We propose that this interaction is confined to specific membrane regions. This assumption brings together the specific biophysical properties of GlyCers (57, 58) with electron microscopic observations made earlier which clearly demonstrated that AFP accumulates at distinct areas at the plasma membrane and cell wall only in the AFP-sensitive fungus *A. niger* but not in the AFP-resistant fungus *P. chrysogenum* (10). These domains might harbor CHSs of classes III and V, which are further known targets of AFP and are exclusively found in filamentous fungi (8). Assuming that unsaturated GlyCers are important for the proper embedding of class III and class V CHSs in these microdomains, a *dtdA* deletion would expose fewer of these CHSs to AFP, being less susceptible. If challenged with AFP, other chitin and glucan synthases could potentially become activated by the cell wall integrity and calcium signaling pathways, thus, fortifying the cell wall stronger in a *dtdA* deletion background than the wild-type background.

**Figure 8.**
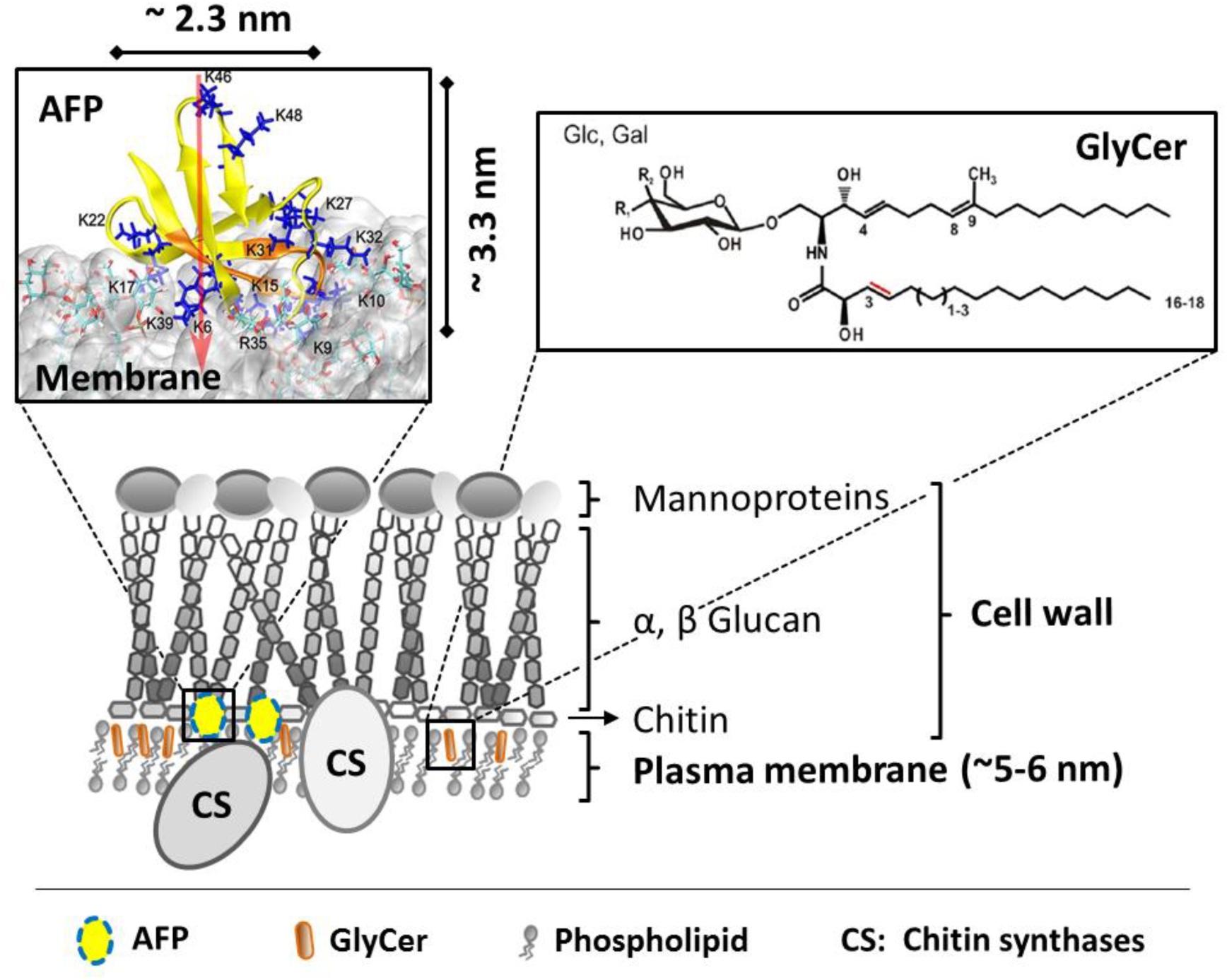
Working model for the mode of action of AFP. See the text for details. Note that the upper left picture is from Utesch et al. (4) (shown with permission from the authors), and displays the MD-simulated interaction of AFP with fungal membrane, including The molecular spatial dimensions of AFP. Yellow: AFP and its N-terminal γ-core are highlighted in orange, respectively. Blue sticks: R and K residues, white cloud: fungal model membrane, red arrow: dipole moments of AFP.

Note that any interaction of AFP with GlyCers remains to be shown; however, the importance of GlyCers for defining the sensitivity of fungi against plant, insect and fungal AMPs have been documented in the literature for a few AMPs already (Suppl. Table S2). This implies that GlyCers could either interact directly with AFP and/or GlyCers are important to embed CHSs in the plasma membrane. It was shown recently that the plant-derived AMP Psd2 interacts preferentially with mimetic membrane domains enriched with GlyCers (60). Previous studies have already shown that a change of membrane fluidity can disrupt membrane protein organization by segregating peripheral and integral proteins and, thereby, interfering with cell wall integrity (61).

In conclusion, this study reports that GlyCers are important determinants for the species-specificity of AFP, suggesting that they can be generally viewed as excellent fungal-specific targets for future antifungal drug development programs. We have revealed that only a minor structural difference in these molecules – the saturation level of their fatty acid chain, controlled by a 2-hydroxy fatty N-acyl-Δ3(*E*)-desaturase – is a key to understanding the inhibitory activity of AFP. Future work will disclose how AFP-mediated inhibition of chitin synthesis is linked with the function of GlyCers for the plasma membrane and, thus, cell wall integrity.

## Supplementary figures

**Figure S1 - Δ3(*E*)-desaturase expression levels at 155 different cultivation conditions of *A. niger*.** Mean transcript levels for An01g09800 are shown based on genome-wide microarray data published in Schäpe et al. (Schäpe P, Kwon MJ, Baumann B, Gutschmann B, Jung S, Lenz S, Nitsche B, Paege N, Schutze T, Cairns TC, Meyer V. 2019. *Updating genome annotation for the microbial cell factory Aspergillus niger using gene co-expression networks*. Nucleic Acids Res 47:559-569).

**Figure S2 - *A. niger* Δ3(*E*)-desaturase transcript expression levels in wild-type and Δ3(*E*)-desaturase mutant strains of filamentous fungi and yeast.** The Ct-values as a measure of the transcript expression levels of Δ3(*E*)-desaturase normalized to that of the actin house-keeping gene analyzed by qRT-PCR are shown. The presence of transcript levels is shown by threshold cycle (Ct)-ratios in the range of around 1–1.2, and their absence, by Ct-ratios in the range of 2–2.6. Data were derived from two independent experiments. Samples of A) were derived from cultures incubated at 30 °C, whereas B) shows samples derived from *A. niger* wild-type strain N402 cultivated at different temperatures to analyze the temperature dependence of Δ3(*E*)-desaturase expression. Desaturase (+) = gene expressed; Desaturase (-) = gene not expressed.

**Figure S3 - Susceptibility of *S. cerevisiae* towards AFP in presence and absence of Δ3(*E*)-desaturase expression.** Growth of *S. cerevisiae* strains BY4741 (wild type) and BBA20.2 (Δ3(*E*)-desaturase knock-in) were tested, according to (Ouedraogo JP, Hagen S, Spielvogel A, Engelhardt S, Meyer V. 2011. *Survival strategies of yeast and filamentous fungi against the antifungal protein AFP.* J Biol Chem 286:13859-68), in the presence of 400 µg/ml AFP or without the peptide. Cultivations were carried out in technical duplicates in microtiter plate format and repeated three times. Error bars express standard deviations. Δ3(*E*)-desaturase (+) = gene expressed; Δ3(*E*)-desaturase (-) = gene not expressed. Data are expressed as mean from two independent experiments each performed in triplicate. Error bars express standard deviations. Δ3(*E*)-desaturase (+) = gene expressed; Δ3(*E*)-desaturase (-) = gene not expressed.

**Figure S4 - MSMS analysis of glycosylceramides in *A. niger* and *P. pastoris*.** MS spectra for the detection of the parental GlyCer ions for *A. niger* N402 18 °C (C) and 37 °C (E), *P. pastoris* BBA21.3 not induced (G) and induced (I). The overlay of the parental ion masses and their fragments are shown: N402 18 °C (D) and 37 °C (F), BBA21.3 not induced (H) and induced (J). R. int. (%) = relative intensity in %; (B) *A. niger dtdA* deletion strain NP1.15. Hex = Hexose

**Figure S5 - Relative chitin and β-1,3-glucan content in wild-type (N402) and *ΔdtdA* deletion strain (NP1.15) of *A. niger*.** The relative amounts of chitin and β-1,3-glucan in % related to control strains, which were set to 100 %, are shown.

## Supplementary tables

**Table S1 -** Primers used in this study. Sequences homologous to the *A. niger* genome are given in upper case, and letters in lower case refer to sequences introduced for PCR/cloning purposes.

**Table S2 - AMPs described to interact with GlyCer.** Protein models were created with SWISS-MODEL (Waterhouse A, Bertoni M, Bienert S, Studer G, Tauriello G, Gumienny R, Heer FT, de Beer TAP, Rempfer C, Bordoli L, Lepore R, Schwede T. 2018. *SWISS-MODEL: homology modelling of protein structures and complexes.* Nucleic Acids Res 46:W296–W303), showing the position of cysteines in yellow.

## Reference List

1. Wiederhold NP. 2017. Antifungal resistance: current trends and future strategies to combat. Infect Drug Resist 10:249–259.

2. Cowen LE. 2008. The evolution of fungal drug resistance: modulating the trajectory from genotype to phenotype. Nat Rev Microbiol 6:187–98.

3. Meyer V. 2008. A small protein that fights fungi: AFP as a new promising antifungal agent of biotechnological value. Appl Microbiol Biotechnol 78:17–28.

4. Utesch T, de Miguel Catalina A, Schattenberg C, Paege N, Schmieder P, Krause E, Miao Y, McCammon JA, Meyer V, Jung S, Mroginski MA. 2018. A computational modeling approach predicts interaction of the antifungal protein AFP from Aspergillus giganteus with fungal membranes via its gamma-core motif. mSphere 3.

5. Barakat H, Spielvogel A, Hassan M, El-Desouky A, El-Mansy H, Rath F, Meyer V, Stahl U. 2010. The antifungal protein AFP from *Aspergillus giganteus* prevents secondary growth of different *Fusarium* species on barley. Appl Microbiol Biotechnol 87:617–624.

6. Paege N, Jung S, Schape P, Muller-Hagen D, Ouedraogo JP, Heiderich C, Jedamzick J, Nitsche BM, van den Hondel CA, Ram AF, Meyer V. 2016. A transcriptome meta-analysis proposes novel biological roles for the antifungal protein AnAFP in *Aspergillus niger*. PLoS One 11:e0165755.

7. Yeaman MR, Yount NY. 2007. Unifying themes in host defence effector polypeptides. Nat Rev Microbiol 5:727–740.

8. Hagen S, Marx F, Ram AF, Meyer V. 2007. The antifungal protein AFP from *Aspergillus giganteus* inhibits chitin synthesis in sensitive fungi. Appl Environ Microbiol 73:2128–2134.

9. Gow NAR, Latge JP, Munro CA. 2017. The fungal cell wall: Structure, biosynthesis, and function. Microbiol Spectr 5.

10. Theis T, Marx F, Salvenmoser W, Stahl U, Meyer V. 2005. New insights into the target site and mode of action of the antifungal protein of *Aspergillus giganteus*. Res Microbiol 156:47–56.

11. Theis T, Wedde M, Meyer V, Stahl U. 2003. The antifungal protein from Aspergillus giganteus causes membrane permeabilization. Antimicrob Agents Chemother 47:588–593.

12. de Vries RP, Riley R, Wiebenga A, Aguilar-Osorio G, Amillis S, Uchima CA, Anderluh G, Asadollahi M, Askin M, Barry K, Battaglia E, Bayram O, Benocci T, Braus-Stromeyer SA, Caldana C, Canovas D, Cerqueira GC, Chen F, Chen W, Choi C, Clum A, Dos Santos RA, Damasio AR, Diallinas G, Emri T, Fekete E, Flipphi M, Freyberg S, Gallo A, Gournas C, Habgood R, Hainaut M, Harispe ML, Henrissat B, Hilden KS, Hope R, Hossain A, Karabika E, Karaffa L, Karanyi Z, Krasevec N, Kuo A, Kusch H, LaButti K, Lagendijk EL, Lapidus A, Levasseur A, Lindquist E, Lipzen A, Logrieco AF, et al. 2017. Comparative genomics reveals high biological diversity and specific adaptations in the industrially and medically important fungal genus *Aspergillus*. Genome Biol 18:28.

13. Ouedraogo JP, Hagen S, Spielvogel A, Engelhardt S, Meyer V. 2011. Survival strategies of yeast and filamentous fungi against the antifungal protein AFP. J Biol Chem 286:13859–13868.

14. Casadevall A, Pirofski LA. 1999. Host-pathogen interactions: redefining the basic concepts of virulence and pathogenicity. Infect Immun 67:3703–3713.

15. Casadevall A, Pirofski LA. 2000. Host-pathogen interactions: basic concepts of microbial commensalism, colonization, infection, and disease. Infect Immun 68:6511–6518.

16. Casadevall A, Pirofski LA. 2003. The damage-response framework of microbial pathogenesis. Nat Rev Microbiol 1:17–24.

17. Leipelt M, Warnecke D, Zähringer U, Ott C, Muller F, Hube B, Heinz E. 2001. Glucosylceramide synthases, a gene family responsible for the biosynthesis of glucosphingolipids in animals, plants, and fungi. J Biol Chem 276:33621–33629.

18. Warnecke D, Heinz E. 2003. Recently discovered functions of glucosylceramides in plants and fungi. Cell Mol Life Sci 60:919–941.

19. Zäuner S, Zähringer U, Lindner B, Warnecke D, Sperling P. 2008. Identification and functional characterization of the 2-hydroxy fatty N-acyl-Delta3(E)-desaturase from Fusarium graminearum. J Biol Chem 283:36734–36742.

20. Leach J, Lang BR, Yoder OC. 1982. Methods for selection of mutants and in vitro culture of *Cochliobolus heterostrophus*. Microbiology 128:1719–1729.

21. Meyer V, Ram AFJ, Punt PJ. 2010. Genetics, genetic manipulation, and approaches to strain improvement of filamentous fungi. Manual of Industrial Microbiology and Biotechnology, Third Edition. American Society of Microbiology.

22. Arentshorst M, Ram AF, Meyer V. 2012. Using non-homologous end-joining-deficient strains for functional gene analyses in filamentous fungi. Methods Mol Biol 835:133–150.

23. Sambrook J, Russell DWDW, Laboratory CSH. 2001. Molecular cloning: a laboratory manual, 3rd ed. Cold Spring Harbor Laboratory.

24. Scorer CA, Clare JJ, McCombie WR, Romanos MA, Sreekrishna K. 1994. Rapid selection using G418 of high copy number transformants of *Pichia pastoris* for high-level foreign gene expression. Biotechnology (N Y) 12:181–184.

25. Becker DM, Guarente L. 1991. High-efficiency transformation of yeast by electroporation. Methods Enzymol 194:182–187.

26. Gouka RJ, van Hartingsveldt W, Bovenberg RA, van Zeijl CM, van den Hondel CA, van Gorcom RF. 1993. Development of a new transformant selection system for *Penicillium chrysogenum*: isolation and characterization of the *P. chrysogenum* acetyl-coenzyme A synthetase gene (*facA*) and its use as a homologous selection marker. Appl Microbiol Biotechnol 38:514–519.

27. Waksman SA, Geiger WB. 1944. The nature of the antibiotic substances produced by *Aspergillus fumigatus*. J Bacteriol 47:391–397.

28. Miedaner T, Reinbrecht C, Schilling AG. 2000. Association among aggressiveness, fungal colonization, and mycotoxin production of 26 isolates of *Fusarium graminearum* in winter rye head blight. J Plant Dis Protect 107:124–134.

29. Cregg JM, Barringer KJ, Hessler AY, Madden KR. 1985. *Pichia pastoris* as a host system for transformations. Mol Cell Biol 5:3376–3385.

30. Winston F, Dollard C, Ricupero-Hovasse SL. 1995. Construction of a set of convenient *Saccharomyces cerevisiae* strains that are isogenic to S288C. Yeast 11:53–55.

31. Rio DC, Ares M, Jr., Hannon GJ, Nilsen TW. 2010. Purification of RNA using TRIzol (TRI reagent). Cold Spring Harb Protoc 2010:pdb prot5439.

32. Cregg JM, Vedvick TS, Raschke WC. 1993. Recent advances in the expression of foreign genes in *Pichia pastoris*. Biotechnology (N Y) 11:905–910.

33. Brachmann CB, Davies A, Cost GJ, Caputo E, Li J, Hieter P, Boeke JD. 1998. Designer deletion strains derived from *Saccharomyces cerevisiae* S288C: A useful set of strains and plasmids for PCR-mediated gene disruption and other applications. Yeast 14:115–132.

34. Kozera B, Rapacz M. 2013. Reference genes in real-time PCR. J Appl Genet 54:391–406.

35. Altschul SF, Gish W, Miller W, Myers EW, Lipman DJ. 1990. Basic local alignment search tool. J Mol Biol 215:403–410.

36. Arentshorst M, Niu J, Ram AFJ. 2015. Efficient generation of *Aspergillus niger* knock-out strains by combining NHEJ mutants and a split marker approach, p 263–272. *In* van den Berg MA, Maruthachalam K (ed), Genetic transformation systems in fungi, Volume 1. Springer International Publishing, Cham.

37. Strohalm M, Hassman M, Kosata B, Kodicek M. 2008. mMass data miner: An open source alternative for mass spectrometric data analysis. Rapid Commun Mass Spectrom 22:905–8.

38. Fortwendel JR, Juvvadi PR, Pinchai N, Perfect BZ, Alspaugh JA, Perfect JR, Steinbach WJ. 2009. Differential effects of inhibiting chitin and 1,3-{beta}-D-glucan synthesis in *ras* and calcineurin mutants of *Aspergillus fumigatus*. Antimicrob Agents Chemother 53:476–482.

39. Ram AF, Arentshorst M, Damveld RA, vanKuyk PA, Klis FM, van den Hondel CA. 2004. The cell wall stress response in *Aspergillus niger* involves increased expression of the glutamine : fructose-6-phosphate amidotransferase-encoding gene (*gfaA*) and increased deposition of chitin in the cell wall. Microbiology 150:3315–3326.

40. Fernandes CM, de Castro PA, Singh A, Fonseca FL, Pereira MD, Vila TV, Atella GC, Rozental S, Savoldi M, Del Poeta M, Goldman GH, Kurtenbach E. 2016. Functional characterization of the *Aspergillus nidulans* glucosylceramide pathway reveals that LCB Delta8-desaturation and C9-methylation are relevant to filamentous growth, lipid raft localization and Psd1 defensin activity. Mol Microbiol 102:488–505.

41. Rittenour WR, Chen M, Cahoon EB, Harris SD. 2011. Control of glucosylceramide production and morphogenesis by the Bar1 ceramide synthase in *Fusarium graminearum*. PLoS One 6:e19385.

42. Levery SB, Momany M, Lindsey R, Toledo MS, Shayman JA, Fuller M, Brooks K, Doong RL, Straus AH, Takahashi HK. 2002. Disruption of the glucosylceramide biosynthetic pathway in *Aspergillus nidulans* and *Aspergillus fumigatus* by inhibitors of UDP-Glc:ceramide glucosyltransferase strongly affects spore germination, cell cycle, and hyphal growth. FEBS Lett 525:59–64.

43. Hillig I, Warnecke D, Heinz E. 2005. An inhibitor of glucosylceramide synthase inhibits the human enzyme, but not enzymes from other organisms. Biosci Biotechnol Biochem 69:1782–5.

44. Kwon MJ, Arentshorst M, Roos ED, van den Hondel CA, Meyer V, Ram AF. 2011. Functional characterization of Rho GTPases in *Aspergillus niger* uncovers conserved and diverged roles of Rho proteins within filamentous fungi. Mol Microbiol 79:1151–1167.

45. Fuchs B, Suss R, Schiller J. 2011. An update of MALDI-TOF mass spectrometry in lipid research. Prog Lipid Res 50:132.

46. Futerman AH, Riezman H. 2005. The ins and outs of sphingolipid synthesis. Trends Cell Biol 15:312–318.

47. . Kusmakow OV. 2006. Versuche zur intrazellulären Lokalisierung der Sterol-Glucosyltransferase und der Glucosylceramid-Synthase in Zellen von *Allium fistulosum L.* Staats- und Universitätsbibliothek Hamburg, Hamburg.

48. Lloyd-Evans E, Pelled D, Riebeling C, Bodennec J, de-Morgan A, Waller H, Schiffmann R, Futerman AH. 2003. Glucosylceramide and glucosylsphingosine modulate calcium mobilization from brain microsomes via different mechanisms. J Biol Chem 278:23594–23599.

49. Sillence DJ, Puri V, Marks DL, Butters TD, Dwek RA, Pagano RE, Platt FM. 2002. Glucosylceramide modulates membrane traffic along the endocytic pathway. J Lipid Res 43:1837–1845.

50. Del Poeta M, Nimrichter L, Rodrigues ML, Luberto C. 2014. Synthesis and biological properties of fungal glucosylceramide. PLoS Pathog 10:e1003832.

51. Huber A, Oemer G, Malanovic N, Lohner K, Kovács L, Salvenmoser W, Zschocke J, Keller MA, Marx F. 2019. Membrane sphingolipids regulate the fitness and antifungal protein susceptibility of *Neurospora crassa*. Front Microbiol 10.

52. Munshi MA, Gardin JM, Singh A, Luberto C, Rieger R, Bouklas T, Fries BC, Del Poeta M. 2018. The role of ceramide synthases in the pathogenicity of *Cryptococcus neoformans*. Cell Rep 22:1392–1400.

53. Dickson RC. 2010. Roles for sphingolipids in *Saccharomyces cerevisiae*. Adv Exp Med Biol 688:21–231.

54. Fernandes CM, Goldman GH, Del Poeta M. 2018. Biological roles played by sphingolipids in dimorphic and filamentous fungi. MBio 9.

55. Singh A, Del Poeta M. 2016. Sphingolipidomics: An important mechanistic tool for studying fungal pathogens. Front Microbiol 7:501.

56. Zhang W, Quinn B, Barnes S, Grabowski GA, Sun Y, Setchell K. 2013. Metabolic profiling and quantification of sphingolipids by liquid chromatography-tandem mass spectrometry. J Glycomics Lipidomics 3:1–8.

57. Gulbins E, Li PL. 2006. Physiological and pathophysiological aspects of ceramide. Am J Physiol Regul Integr Comp Physiol 290:R11–26.

58. Sonnino S, Prinetti A. 2013. Membrane domains and the “lipid raft” concept. Curr Med Chem 20:4–21.

59. Briolay A, Bouzenzana J, Guichardant M, Deshayes C, Sindt N, Bessueille L, Bulone V. 2009. Cell wall polysaccharide synthases are located in detergent-resistant membrane microdomains in oomycetes. Appl Environ Microbiol 75:1938–1949.

60. Amaral VSG, Fernandes CM, Felicio MR, Valle AS, Quintana PG, Almeida CC, Barreto-Bergter E, Goncalves S, Santos NC, Kurtenbach E. 2019. Psd2 pea defensin shows a preference for mimetic membrane rafts enriched with glucosylceramide and ergosterol. Biochim Biophys Acta Biomembr 1861:713–728.

61. Scheinpflug K, Wenzel M, Krylova O, Bandow JE, Dathe M, Strahl H. 2017. Antimicrobial peptide cWFW kills by combining lipid phase separation with autolysis. Sci Rep 7:44332.

62. Waterhouse A, Bertoni M, Bienert S, Studer G, Tauriello G, Gumienny R, Heer FT, de Beer TAP, Rempfer C, Bordoli L, Lepore R, Schwede T. 2018. SWISS-MODEL: homology modelling of protein structures and complexes. Nucleic Acids Res 46:W296–W303.

63. Aerts AM, Francois IE, Meert EM, Li QT, Cammue BP, Thevissen K. 2007. The antifungal activity of RsAFP2, a plant defensin from *Raphanus sativus*, involves the induction of reactive oxygen species in *Candida albicans*. J Mol Microbiol Biotechnol 13:243–247.

64. Thevissen K, Cammue BP, Lemaire K, Winderickx J, Dickson RC, Lester RL, Ferket KK, Van Even F, Parret AH, Broekaert WF. 2000. A gene encoding a sphingolipid biosynthesis enzyme determines the sensitivity of *Saccharomyces cerevisiae* to an antifungal plant defensin from dahlia (*Dahlia merckii*). Proc Natl Acad Sci U S A 97:9531–9536.

65. Thevissen K, Warnecke DC, Francois IE, Leipelt M, Heinz E, Ott C, Zähringer U, Thomma BP, Ferket KK, Cammue BP. 2004. Defensins from insects and plants interact with fungal glucosylceramides. J Biol Chem 279:3900–3905.

66. Ramamoorthy V, Cahoon EB, Li J, Thokala M, Minto RE, Shah DM. 2007. Glucosylceramide synthase is essential for alfalfa defensin-mediated growth inhibition but not for pathogenicity of *Fusarium graminearum*. Mol Microbiol 66:771–786.

