## Supplemental Figure 1 for "Species-specific differences in the susceptibility of fungi towards the antifungal protein AFP depend on C3 saturation of glycosylceramides"

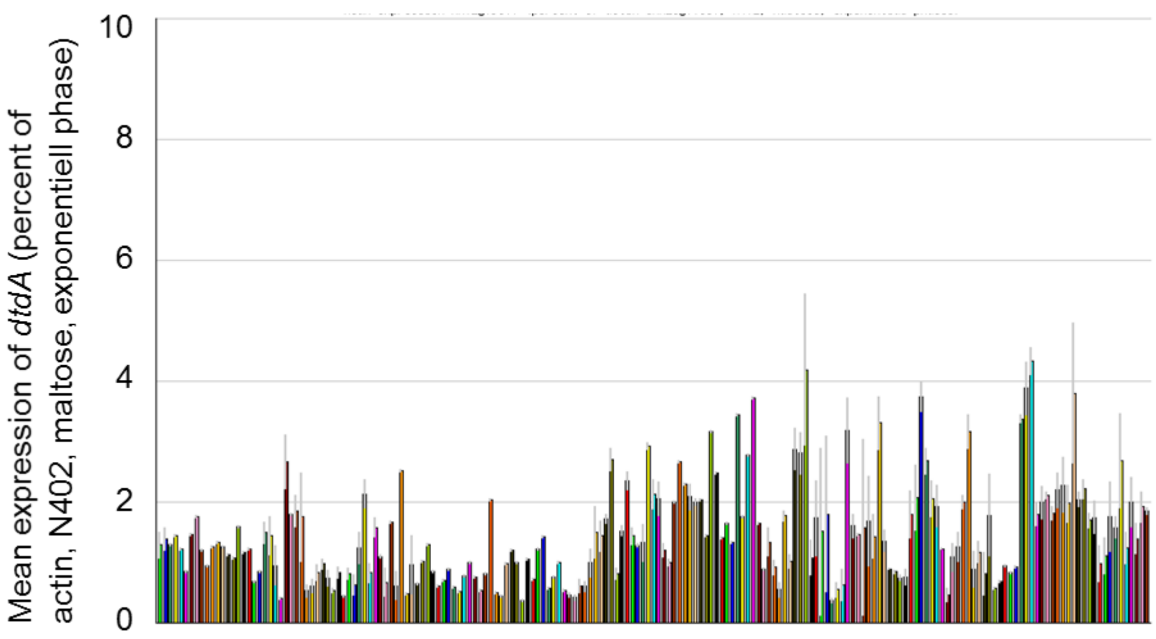


**Figure S1 - Δ3(E)-desaturase expression levels at 155 different cultivation conditions of *A. niger*.** Mean transcript levels for An01g09800 are shown based on genome-wide microarray data published in Schäpe et al. (Schäpe P, Kwon MJ, Baumann B, Gutschmann B, Jung S, Lenz S, Nitsche B, Paege N, Schutze T, Cairns TC, Meyer V. 2019. Updating genome annotation for the microbial cell factory *Aspergillus niger* using gene co-expression networks. Nucleic Acids Res 47:559–569).
