## Supplemental Figure 2 for "Species-specific differences in the susceptibility of fungi towards the antifungal protein AFP depend on C3 saturation of glycosylceramides"

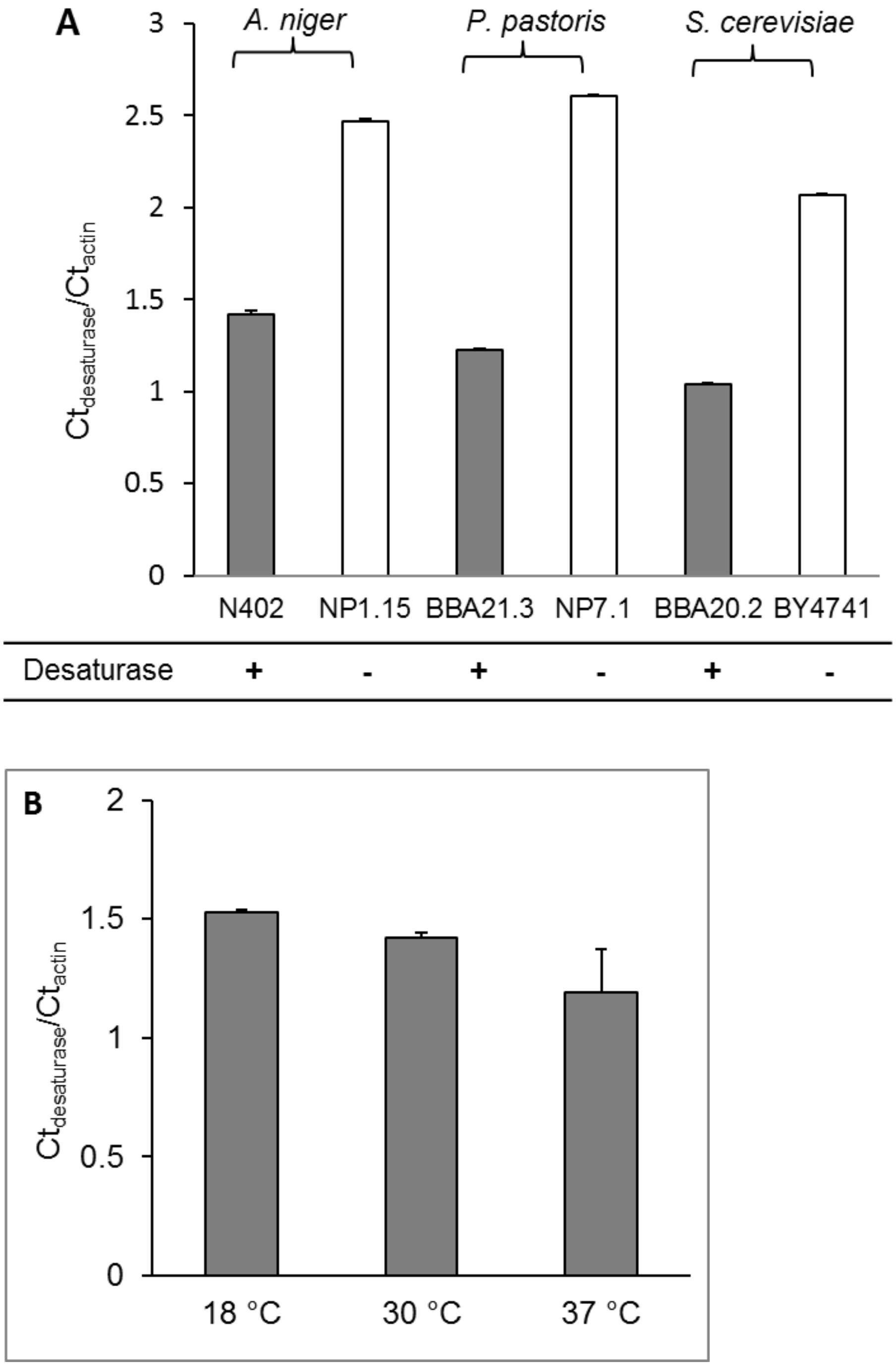


**Figure S1 - *A. niger* Δ3(E)-desaturase transcript expression levels in wild-type and Δ3(E)-desaturase mutant strains of filamentous fungi and yeast.** The Ct-values are shown as a measure of the transcript expression levels of Δ3(E)-desaturase normalized to that of the actin house-keeping gene analyzed by qRT-PCR. The presence of transcript levels is shown by threshold cycle (Ct)-ratios in the range of around 1–1.2, their absence, by Ct-ratios in the range of 2–2.6. Data were derived from two independent experiments. Samples of A) were derived from cultures incubated at 30 °C, whereas B) shows samples derived from *A. niger* wild-type strain N402 cultivated at different temperatures to analyze temperature dependence of $\Delta$3(E)-desaturase expression. Desaturase (+) = gene expressed; Desaturase (-) = gene not expressed.
