## Supplemental Figure 3 for "Species-specific differences in the susceptibility of fungi towards the antifungal protein AFP depend on C3 saturation of glycosylceramides"

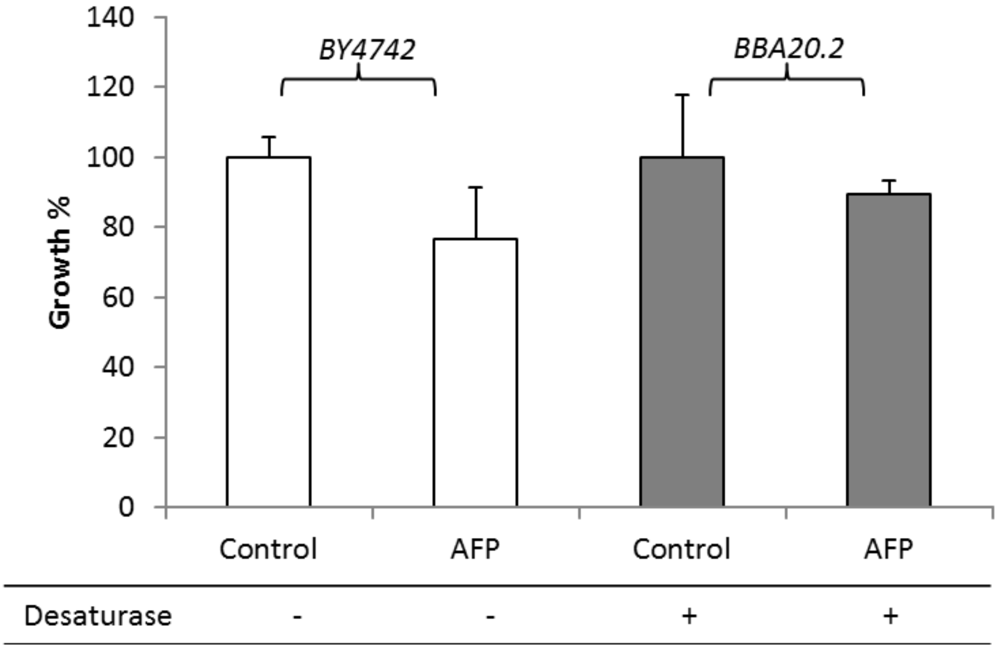


**Figure S3 - Susceptibility of *S. cerevisiae* towards AFP in the presence and absence of Δ3(E)-desaturase expression.** Growth of *S. cerevisiae* strains BY4741 (wild type) and BBA20.2 (Δ3(E)-desaturase knock-in) were tested, according to (Ouedraogo JP, Hagen S, Spielvogel A, Engelhardt S, Meyer V. 2011. Survival strategies of yeast and filamentous fungi against the antifungal protein AFP. J Biol Chem 286:13859–13868.), in the presence of 400 µg/ml AFP or without the peptide. Cultivations were carried out in technical duplicates in microtiter plate format and repeated three times. Error bars express standard deviations. Δ3(E)-desaturase (+) = gene expressed; Δ3(E)-desaturase (-) = gene not expressed. Data are expressed as mean from two independent experiments, each performed in triplicate. Error bars express standard deviations. Δ3(E)-desaturase (+) = gene expressed; Δ3(E)-Desaturase (-) = gene not expressed.
