## Supplemental Figure 4 for "Species-specific differences in the susceptibility of fungi towards the antifungal protein AFP depend on C3 saturation of glycosylceramides"

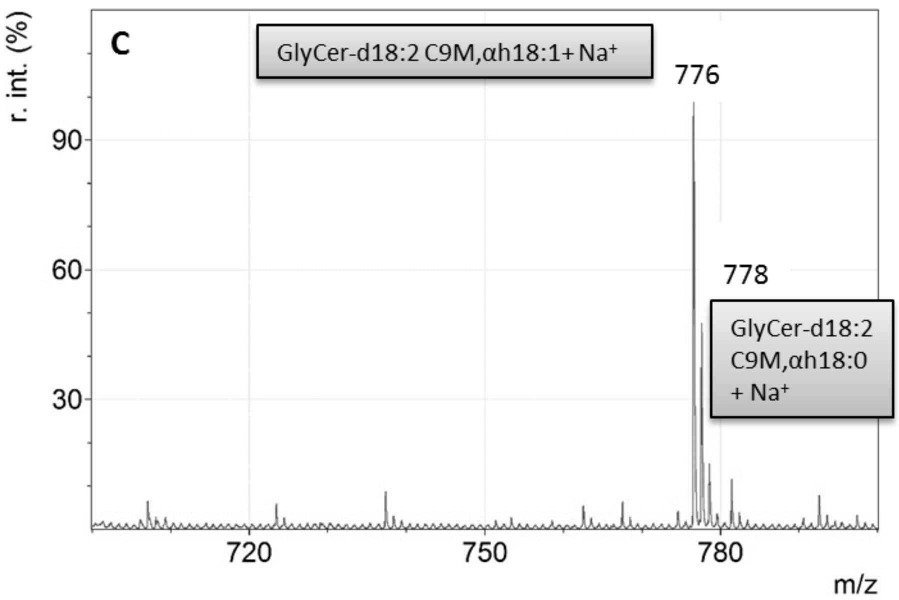

C
GlyCer-d18:2 C9M,αh18:1+ Na+
776
778
GlyCer-d18:2
C9M,αh18:0
+ Na+
r. int. (%)
90
60
30
720
750
780
m/z


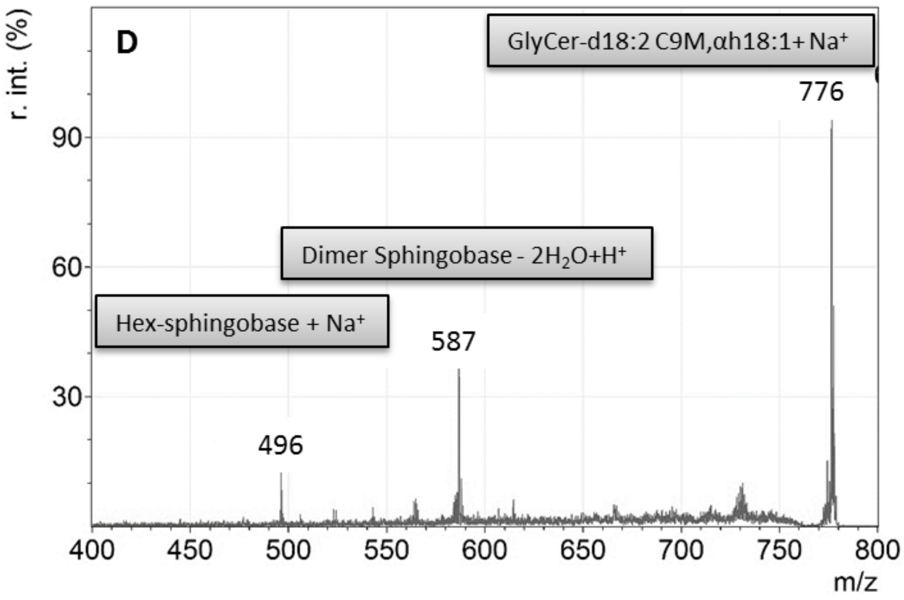

D
GlyCer-d18:2 C9M,αh18:1+ Na+
776
Dimer Sphingobase - 2H2O+H+
Hex-sphingobase + Na+
587
496
r. int. (%)
90
60
30
400
450
500
550
600
650
700
750
800
m/z

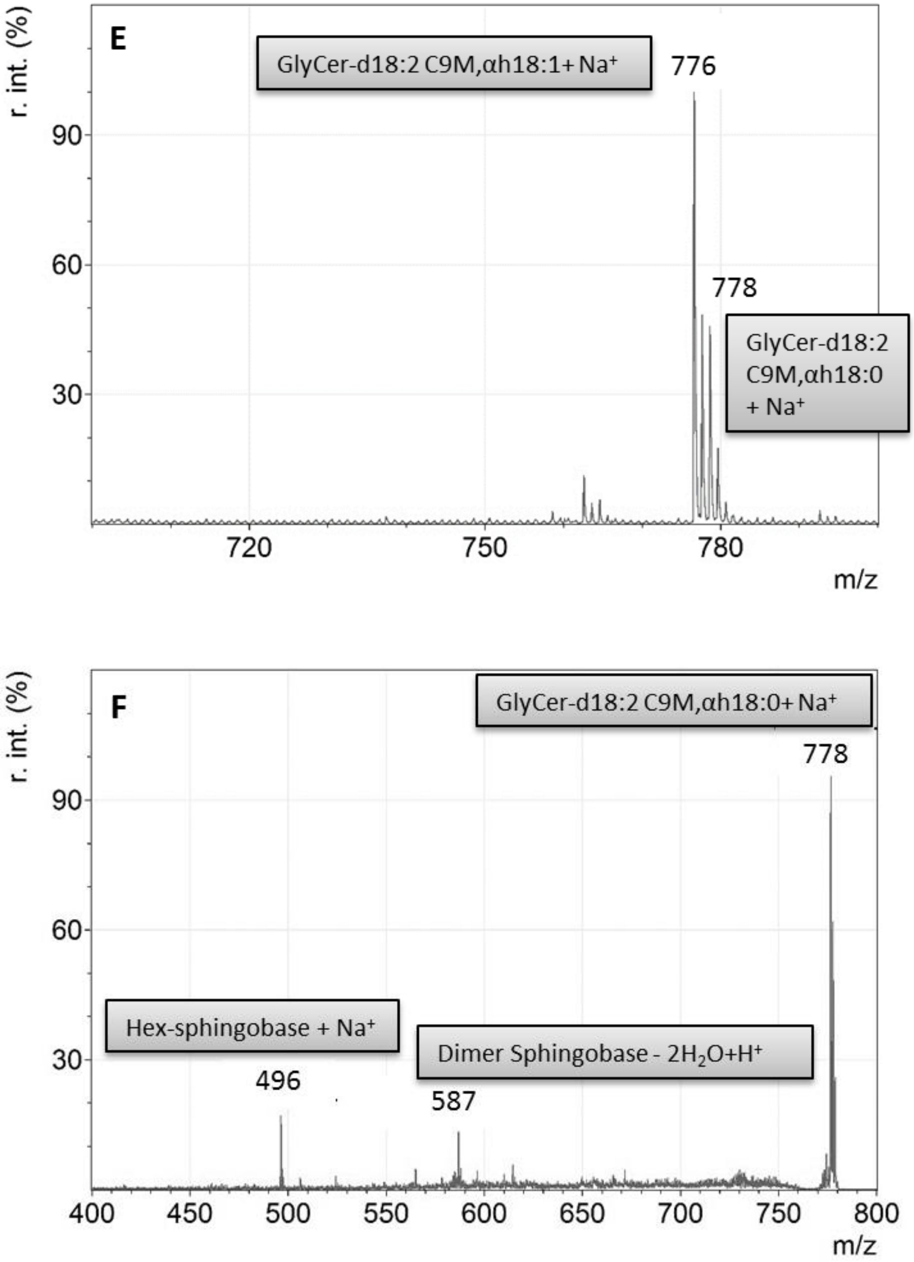

E
GlyCer-d18:2 C9M,αh18:1+ Na+
776
778
GlyCer-d18:2 C9M,αh18:0 + Na+
r. int. (%)
90
60
30
720
750
780
m/z
F
GlyCer-d18:2 C9M,αh18:0+ Na+
778
Hex-sphingobase + Na+
496
Dimer Sphingobase - 2H2O+H+
587
r. int. (%)
90
60
30
400
450
500
550
600
650
700
750
800
m/z

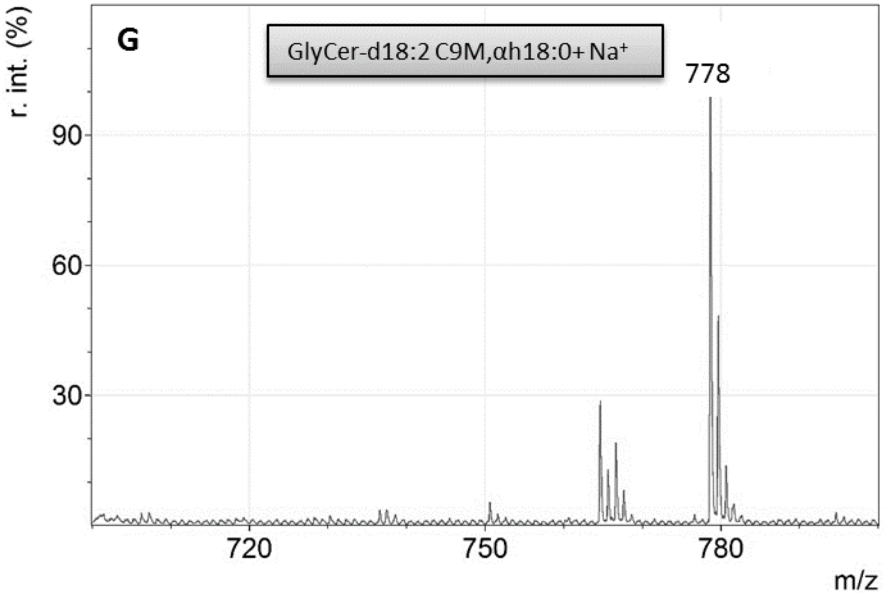
G
GlyCer-d18:2 C9M,αh18:0+ Na+
778
r. int. (%)
90
60
30
720
750
780
m/z

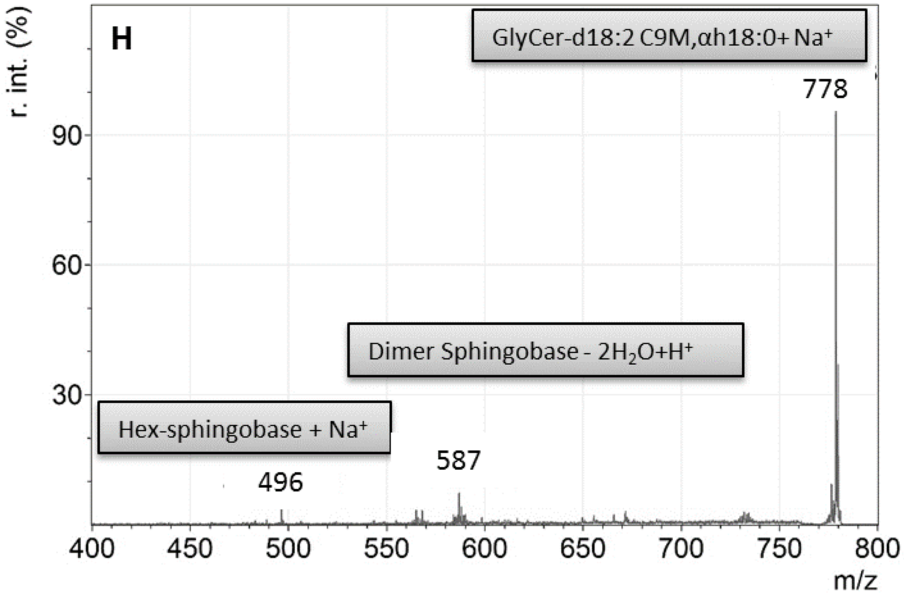
H
GlyCer-d18:2 C9M,αh18:0+ Na+
778
Dimer Sphingobase - 2H2O+H+
Hex-sphingobase + Na+
496
587
r. int. (%)
90
60
30
400
450
500
550
600
650
700
750
800
m/z

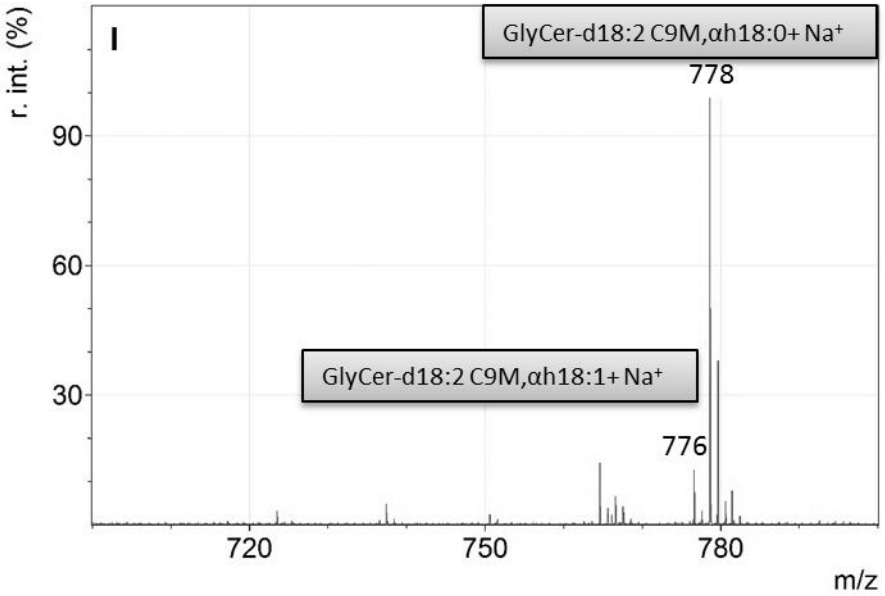


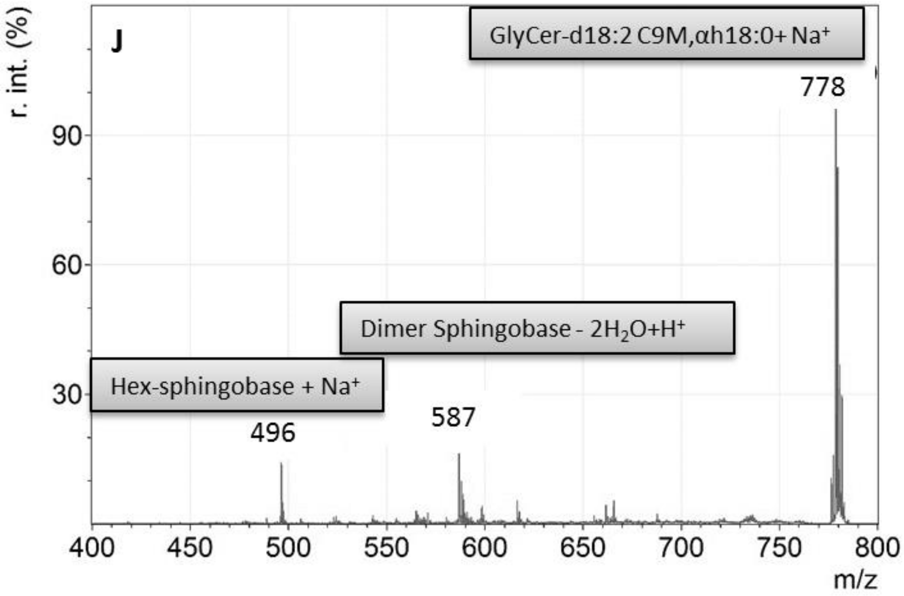


**Figure S4 - MSMS analysis of glycosylceramides in *A. niger* and *P. pastoris*.** MS spectra for the detection of the parental GlyCer ions for *A. niger* N402 18 °C (C) and 37 °C (E), *P. pastoris* BBA21.3 not induced (G) and induced (I). The overlay of the parental ion masses and their fragments are shown: N402 18 °C (D) and 37 °C (F), BBA21.3 not induced (H) and induced (J). R. int. (%) = relative Intensity in %; (B) *A. niger dtdA* deletion strain NP1.15. Hex = Hexose
