## Supplemental Figure 5 for "Species-specific differences in the susceptibility of fungi towards the antifungal protein AFP depend on C3 saturation of glycosylceramides"

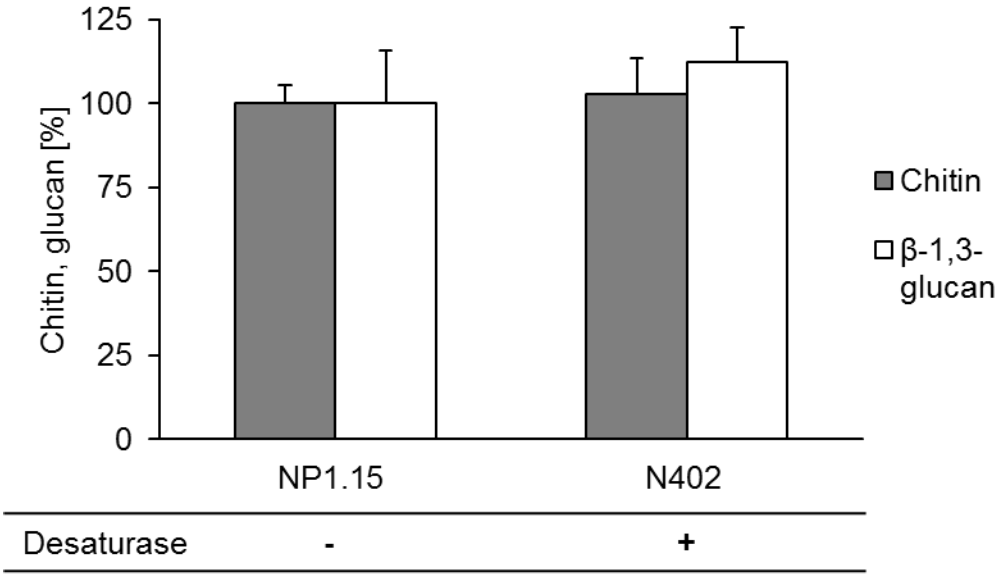


**Figure S5 - Relative chitin and β-1,3-glucan content in wild-type (N402) and *ΔdtdA* deletion strain (NP1.15) of *A. niger***. The relative amounts of chitin and β-1,3-glucan in % related to control strains, which were set to 100 %, are shown.
