## Supplemental Table 1 for "Species-specific differences in the susceptibility of fungi towards the antifungal protein AFP depend on C3 saturation of glycosylceramides"

**Table S1 -** Primers used in this study. Sequences homologous to the *A. niger* genome are given in upper case, and letters in lower case refer to sequences introduced for PCR/cloning purposes.

| Primer Code | Localization | Orientation | Sequence |
|---|---|---|---|
| 459 | 5`end *pyrG* region | fw | aagccgctgctggaattgTGTAAAACGACGGCCAGT |
| 460 | 3`end *pyrG* region | rev | CGATGGATAATTGTGCCGTGT |
| 461 | 5`end *pyrG* | fw | ATTGACCTACAGCGCACGC |
| 462 | In *pyrG* | rev | CCGGTAGCCAAAGATCCCTT |
| 677 | Upstream *An01g09800* | fw | ACACATGGGAGGGGTAATGA |
| 678 | Upstream *An01g09800* with *AopyrG* start | rev | tcactggccgtcgttttacaCATGTTATCCGACCCATCCC |
| 679 | Downstream *An01g09800* with *AopyrG* end | fw | acacggcacaattatccatcgTCTCACTAGCAGACGAGGTT |
| 680 | Downstream | rev | ACCCATCGATTAGGGAGAGG |

| | | | |
|---|---|---|---|
| | *An01g09800* | | |
| 716 | Downstream 680 | rev | TCACCACCACAATCTACCCC |
| 936 | pYip5 with 5` site *tef* | fw | gctgatgagctttaccgcagCAAAATGTTTCTACTCCTTT |
| 937 | 3`site *tef* with 5`site Kozak sequence Yeast and *An01g09800* | | gggtccatggtggcggcgggCTTAGATTAGATTGCTATGC |
| 938 | *An01g09800* with 3`end *tef* Kozak sequence yeast | fw | aatctaagcccgccgccaccATGGACCCTTCCACCTTTATexon |
| 939 | 3`site *An01g09800* with 5`end *cyc* | rev | tgacataactaattacatgaCTAAACTTCCGCTTGATTTC |
| 940 | 5`end *cyc* with 3`end *An01g09800* | fw | gaaatcaagcggaagtttagTCATGTAATTAGTTATGTCA |
| 941 | 3` end *cyc* | rev | caccgaaacgcgcgaggcagAGCGTCCCAAAACCTTCTCA |

| | | | |
|---|---|---|---|
| | with pYIp5 | | |
| 1436 | 5`end *An01g09800* with BamHI site | fw | gatccacataATGGACCCTTCCACCTTTAT |
| 1439 | *5`An01g09800* | fw | ATGGACCCTTCCACCTTTAT |
| 1440 | *3`An01g09800* | rev | TCAGCTAAACTTCCGCTTGA |
| 1443 | *3`An01g09800 with* EcoRI site | rev | aattcTCAGCTAAACTTCCGCTTG |
| 1529 | *3`An01g09800 with* NotI site | rev | gcgaattaattcgcggccgcTCAGCTAAACTTCCGCTTGA |
| 1530 | *5`An01g09800 with* EcoRI | fw | gatcctacgtagaattcattATGGACCCTTCCACCTTTAT |
