## Supplemental Table 2 for "Species-specific differences in the susceptibility of fungi towards the antifungal protein AFP depend on C3 saturation of glycosylceramides"

**Table S2 - AMPs described to interact with GlyCer.** Protein models were created with SWISS-MODEL (Waterhouse A, Bertoni M, Bienert S, Studer G, Tauriello G, Gumienny R, Heer FT, de Beer TAP, Rempfer C, Bordoli L, Lepore R, Schwede T. 2018. *SWISS-MODEL: homology modelling of protein structures and complexes.* Nucleic Acids Res 46:W296–W303.), showing the position of cysteines in yellow.

| **Name** | **Organism** | **Structure** | **Uniprot-no.** | **Source** |
|---|---|---|---|---|
| RsAFP | *Raphanus sativus* | 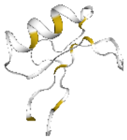 | P69241 | (63) |
| Psd1 | *Pisum sativum* | 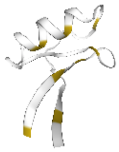 | P81929 | (40), |
| Psd2 | *Pisum sativum* | 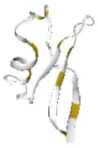 | P81930 | (60), |
| DmAMP1 | *Dahlia merckii* | 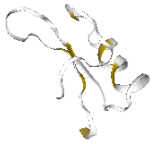 | P0C8Y4 | (64), |
| Heliomicin | *Heliothis virescens* | 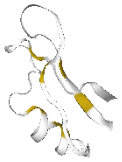 | P81544 | (65), |
| *Ms*Def1 | *Medicago sativa* | 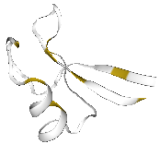 | Q9FPM3 | (66), |

| | | | | |
|---|---|---|---|---|
| PAF | *P. chrysogenum* | 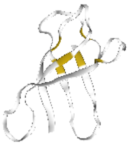 | D0EXD3 | (51), |
| AFP | *A. giganteus* | 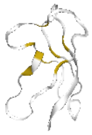 | P17737 | this work |
